# CCR2 silencing in sensory neurons blocks bone cancer progression

**DOI:** 10.1101/2024.05.29.596531

**Authors:** Élora Midavaine, Jérôme Côté, Alexandra Trépanier, Sakeen W. Kashem, Marc-André Dansereau, Jean-Michel Longpré, Martine Charbonneau, Claire Dubois, Ashley M. Jacobi, Scott D. Rose, Mark A. Behlke, Philippe Sarret

## Abstract

The peripheral nervous system contributes to cancer growth, in part by shaping the immunological niche of the tumor. How the nervous system influences bone cancer progression, and whether the underlying neuroimmune pathways can be targeted therapeutically, remain unclear. Here we demonstrate a profound influence of the peripheral nervous system on tumor progression that can be countered by silencing chemokine receptor signaling in sensory neurons. Axotomy of the tumor-innervating femoral nerve inhibits tumor progression in animals bearing bone cancer, whereas intrathecal delivery of the tumor-associated proinflammatory chemokine CCL2 promotes both tumor growth and allodynia. Silencing CCR2 in dorsal root ganglion (DRG) neurons with a newly developed lipid nanoparticle-formulated Dicer-substrate siRNA impedes tumor progression and pathological bone remodeling, and relieves bone cancer-associated pain. Mechanistically, bone cancer drives CCR2-dependent transport of substance P and CGRP along the tumor-innervating femoral nerve, and these neuropeptides expand the tumor-associated macrophage population; silencing CCR2 in DRG neurons normalizes the neuropeptide milieu and ameliorates altered bone remodeling. We thus define a targetable neuroimmune axis that contributes to cancer progression.

**Highlights:**

- Cancer progression activates sensory neurons, driving pain hypersensitivity and neuropeptide release.
- Axotomy of the tumor-innervating femoral nerve impedes tumor progression.
- CCL2–CCR2 signaling in DRG neurons promotes pain hypersensitivity and cancer growth.
- Silencing CCR2 in the DRG reduces pain hypersensitivity, tumor-associated macrophage numbers and cancer growth.

## Introduction

Bone metastases cause intractable pain and carry a very unfavorable prognosis. Bone is the most frequent site of distant relapse in breast cancer, and up to 70% of patients with advanced breast or prostate cancer develop skeletal metastases^1^. Once seeded in bone, these tumors are incurable and ultimately lethal, and patient management is compromised by the limited efficacy of currently available analgesics^2^. Anti-osteolytic agents combined with radiotherapy and chemotherapy remain the mainstay of treatment, but their anticancer efficacy is limited^3^.

Clinical and preclinical work has identified a communication network between the peripheral nervous system and cancer that includes neuro-neoplastic synapses contributing to tumor growth^4,5,6,7,8,9^. Sympathetic and sensory fibers are found within tumor beds^10^, and many stromal and cancer cells express receptors for neuropeptides^11,12^. These neuromediators influence tumor progression by several routes: direct mitogenic effects on cancer cells^13^, modulation of the desmoplastic stroma, a shift in infiltrating immune cells toward a tumor-promoting phenotype, enhanced angiogenesis^7^ and promotion of metastasis^14^.

The CCR2 receptor and its ligand CCL2 are well-established drivers of neuroinflammation, tumor growth and cancer-associated pain^15,16,17,18^. CCL2 levels correlate with poor treatment response and early mortality in patients with metastases^19,20,21^, and high CCL2 in the tumor environment is closely associated with the recruitment of CCR2^+^ monocytes to the tumor niche, which increases drug resistance and accelerates malignant progression^17,22,23,24^.

Given the intimate relationship between inflammation and metastatic growth, we set out to define the link between metastasis-induced neuroinflammation and tumor progression. We show that bone cancer increases CCR2 expression and neuronal activation in dorsal root ganglion (DRG) neurons, which in turn drives antidromic transport of substance P and CGRP into the tumor milieu. By expanding the macrophage population, these neuromodulators promote tumor angiogenesis and pathological bone remodeling, resulting in enhanced tumor growth. Consistent with a tumor-promoting role for neuron-derived factors released into the tumor niche, axotomy significantly diminished bone cancer growth. Intrathecal delivery of CCL2 induced pain sensitivity, neuropeptide production and tumor progression, whereas silencing CCR2 within the DRG reduced bone cancer pain, neuropeptide transport, tumor-associated macrophage numbers and, consequently, tumor progression. Together, these findings define a cancer-associated neuroimmune mechanism that can be selectively targeted to limit bone cancer progression.

## Results

### Innervation of femoral bone tumors contributes to tumor growth

Sensory and autonomic neurons contribute to tumor growth and are often intimately associated with the tumor microenvironment. We therefore asked how sensory innervation contributes to the progression of tumors growing within bone. The femur is innervated principally by the femoral nerve, which carries both sensory and motor fibers. To determine whether local neurogenic inflammation promotes tumor progression, we axotomized the femoral nerve in rats (Fig. 1A and B). A two-month recovery period was imposed so that surgical inflammation could resolve and would not itself influence tumor progression. Rats then underwent surgical implantation of syngeneic MRMT-1 breast carcinoma cells into the medullary cavity of the femur. Twenty days after implantation, axotomized animals had significantly lower tumor weights than rats subjected to the sham axotomy procedure (Fig. 1C and D).

**Figure 1.**
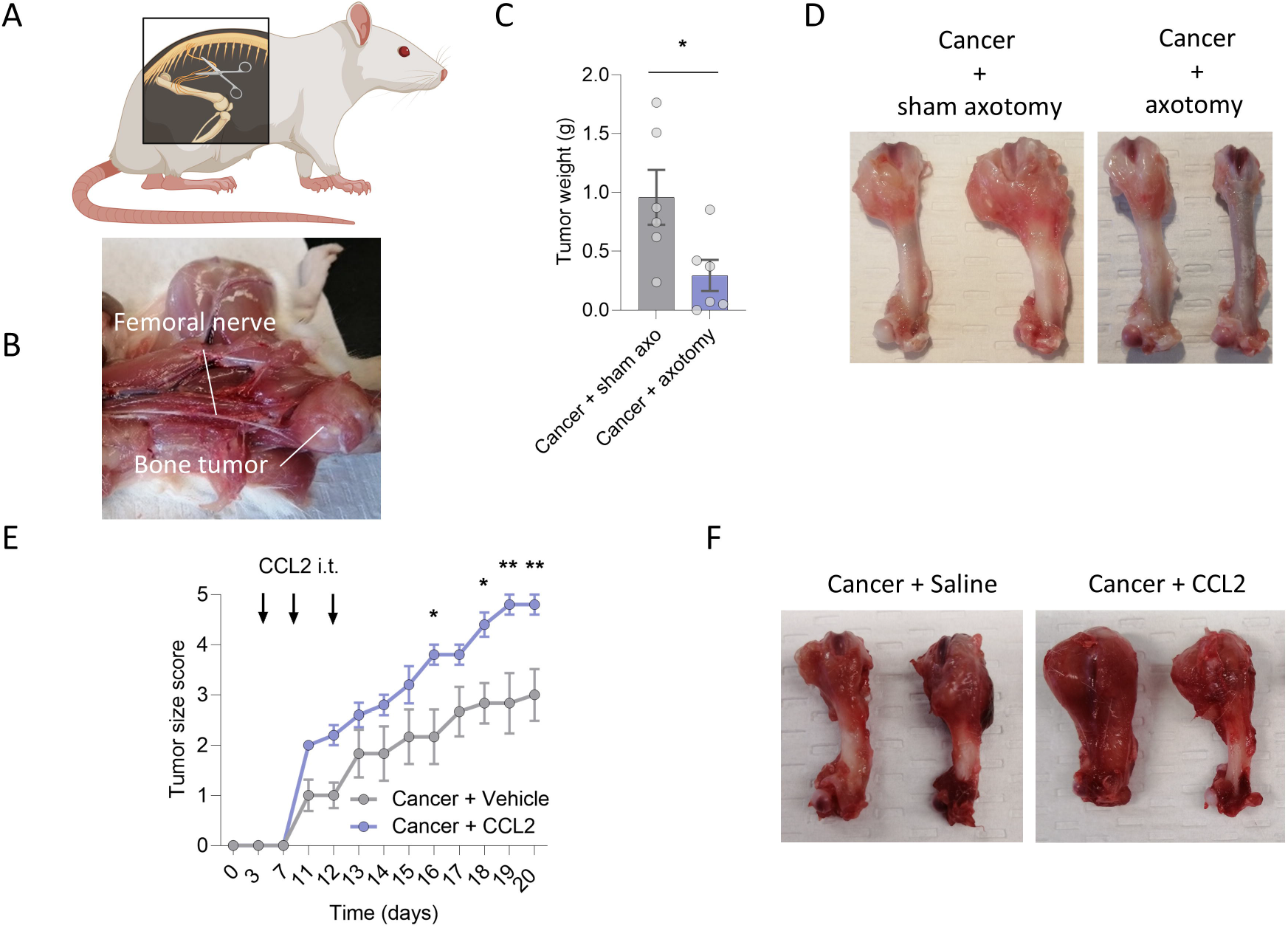
Innervation of the femur contributes to tumor growth. (A–B) Photograph showing tumor innervation by the femoral nerve, and schematic representation of the denervation procedure. (C–D) Surgical denervation of the femoral nerve prior to cancer implantation significantly reduced tumor size at POD 20. Sham axo = sham axotomy. (E) Repeated intrathecal injection of CCL2 accelerated tumor growth. (F) Representative photographs of saline- and CCL2-injected cancer-bearing animals. Data are presented as mean ± SEM. Mann–Whitney test in (C) and two-way ANOVA followed by Sidak’s multiple comparisons test in (E). *P < 0.05, **P < 0.01.

We reasoned that, if neurogenic inflammation arising from the femoral nerve promotes tumor growth, repeated intrathecal injection of a known cancer-associated proinflammatory agent should have the opposite effect. Intrathecally delivered compounds readily reach the DRG, which contains the cell bodies of sensory neurons. CCL2 drives neuronal excitability^25^, and CCR2 blockade limits substance P transport to the periphery in a model of chronic inflammatory pain induced by intraplantar CFA^26^. Consistent with our hypothesis, tumor-bearing rats given exogenous intrathecal CCL2 exhibited significantly greater tumor growth than saline-injected animals (Fig. 1E and F, Supplementary Fig. 1). These results demonstrate that the peripheral nervous system exerts a significant influence on tumor progression.

### Silencing CCR2 in the DRG reduces tumor growth and cancer-induced pain behaviors

We next sought a therapeutic approach to counter the contribution of tumor innervation to tumoral expansion. CCR2 is expressed in sensory neuron cell bodies within the DRG and in the central terminals of these cells in the dorsal horn of the spinal cord, and its expression is upregulated in cancer (Fig. 2A and B, Supplementary Fig. 2D). To interrogate the role of CCR2 expressed on sensory neurons, we tested the ability of two Dicer-substrate small interfering RNAs (DsiRNAs) to downregulate CCR2. We first compared the silencing efficacy of two CCR2-targeting sequences, each with two methylation patterns, in CCR2-expressing HEK cells (Supplementary Tables 1 and 2). On the basis of its silencing efficacy, the CCR2-6 sequence was selected over CCR2-5 for all subsequent experiments (Supplementary Fig. 2A). The methylation patterns tested did not compromise silencing efficacy and are known to confer resistance to degradation and to reduce immunogenicity, thereby increasing potency and limiting side effects^27^. The methylated CCR2-6 sequence and its non-targeting methylated control (NC5) were encapsulated in Neuro9 lipid nanoparticles (LNP-DsiRNA) using a microfluidic device to ensure effective cell penetration (Fig. 2B).

**Figure 2.**
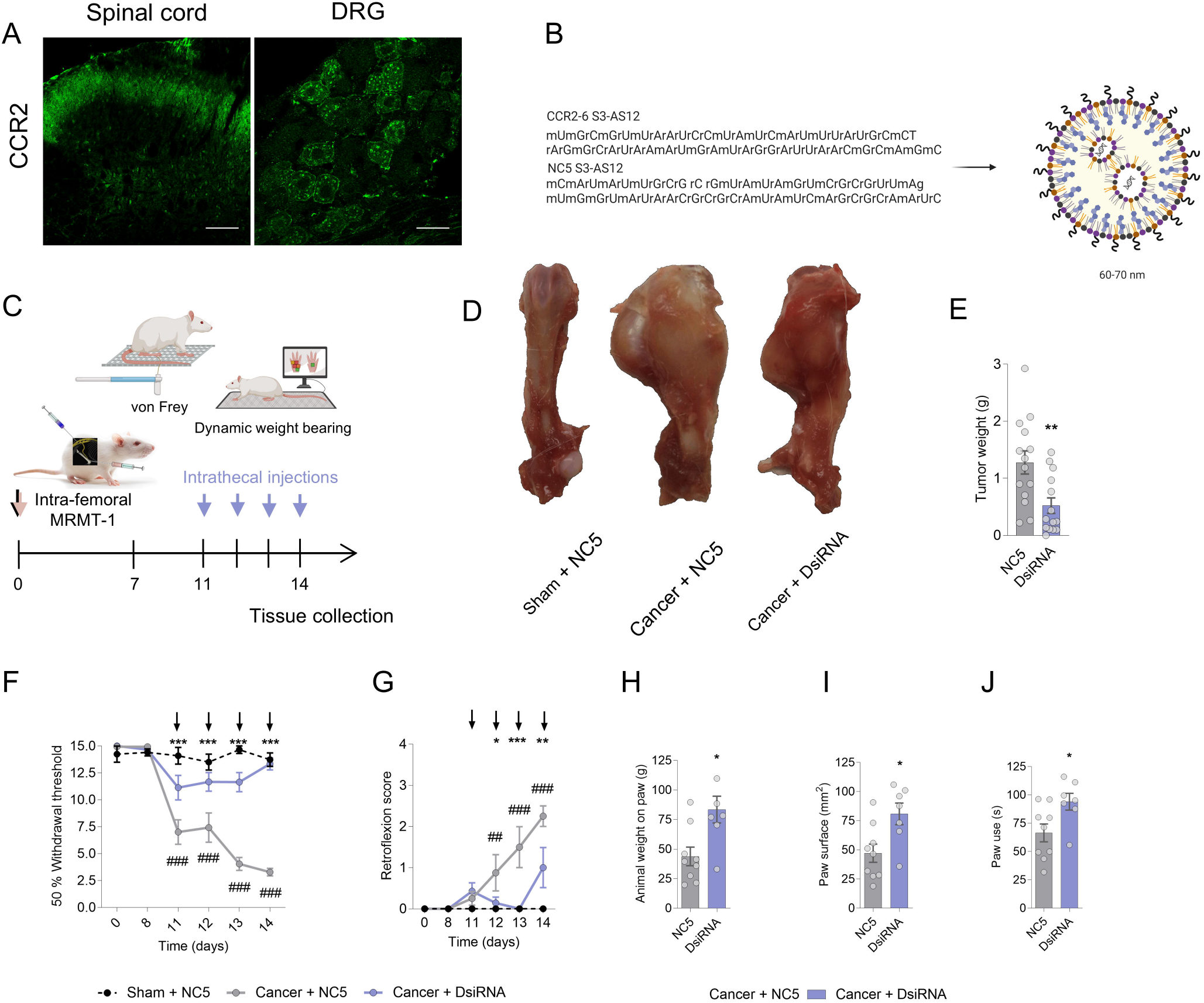
Silencing CCR2 expression in the DRG reduces tumor burden in the femur and tumor-induced pain hypersensitivity. CCR2 is expressed in the somatosensory pathway, including (A) the spinal cord and (B) the DRG. Scale bars, 100 µm and 20 µm. (C) Schematic representation of the study design and of the behavioral tests used to measure stimulus-evoked and non-evoked pain-related behaviors in tumor-bearing rats. (D) Representative photographs of the ipsilateral femurs of NC5-treated shams and of NC5- and DsiRNA-treated cancer-implanted animals at POD 14. (E) Tumor weights at POD 14 in NC5- and DsiRNA-treated animals (cancer+NC5, n = 14; cancer+DsiRNA, n = 14; LNP-DsiRNA, 297 pmol/rat, i.t.). (F–G) Mechanical von Frey thresholds and tumor-bearing paw retroflexion scores used to evaluate the efficacy of repeated daily administration of LNP-DsiRNA (297 pmol/rat, i.t.) between POD 11 and POD 14. LNP-DsiRNA efficacy was also assessed with a dynamic weight-bearing device to measure (H) the weight applied to the ipsilateral hind paw, (I) the paw surface against the pressure pad and (J) paw use. Data are presented as mean ± SEM. Mann–Whitney test in (E, H–J) and two-way ANOVA followed by Sidak’s multiple comparisons test in (F–G). *P < 0.05, **P < 0.01, ***P < 0.001. # compared with the sham+NC5 group and * compared with the cancer+NC5 group.

To assess whether these DsiRNAs modulate CCR2-induced signaling *in vivo*, we first asked whether they reduce mechanical hypersensitivity in a CCL2-induced acute pain model, as previously described^28^. Spinal administration of LNP-encapsulated anti-CCR2 DsiRNA (LNP-DsiRNA, 297 pmol/rat, i.t.) once daily for the two days preceding CCL2 delivery prevented CCL2-induced mechanical allodynia, confirming the silencing efficacy of our LNP-DsiRNA (Supplementary Fig. 2B and C).

We next implanted MRMT-1 breast carcinoma cells into the medullary cavity of the femur and performed daily intrathecal injections of anti-CCR2 LNP-DsiRNA (297 pmol/rat) from day 11 to day 14 after implantation. Compared with NC5, repeated LNP-DsiRNA injections produced a striking reduction in tumor growth in cancer-bearing animals (Fig. 2C–E).

Because the clinical management of chronic bone cancer pain remains challenging given its complex and incompletely understood etiology, we next asked whether LNP-DsiRNA is analgesic in this setting. Chronic administration of LNP-DsiRNA reversed mechanical hypersensitivity and spontaneous paw retroflexion in cancer-bearing animals (Fig. 2F and G), and increased weight bearing, the paw surface applied by the affected limb and the time spent using that limb in freely moving tumor-bearing rats (Fig. 2H–J). Importantly, LNP-DsiRNA altered neither bone marrow metabolism, bone density nor bone vascularization in the tumor-free contralateral femur, and did not affect weight gain, indicating that repeated treatment was well tolerated (Supplementary Fig. 3A–D). Repeated administration of anti-CCR2 DsiRNA also limited the cancer-induced increases in CCR2 expression and neuronal activation in the DRG (Supplementary Fig. 2D and E) and reduced the expression of substance P and CGRP that accompanies the development of bone cancer pain, further confirming target engagement and the analgesic action of CCR2 knockdown (Supplementary Fig. 4A and B).

To determine whether the antitumor effect is simply a consequence of the antinociceptive activity of LNP-DsiRNA, we tested whether ziconotide (Prialt) also inhibits tumor progression. Ziconotide is an N-type calcium channel blocker used intrathecally in palliative therapy for patients with cancer pain^29^. Administered twice daily from day 11 to day 14 after cancer cell implantation, ziconotide significantly reversed bone cancer pain but had no tumor-modulating effect (Supplementary Fig. 5A–D). Analgesia alone is therefore not sufficient to limit tumor growth in this model. Overall, targeting CCR2 expression in the intrathecal compartment produces profound tumor-limiting and analgesic effects.

### Increased substance P and CGRP transport into the tumor-innervating femoral nerve is reduced following CCR2 downregulation

To investigate how intrathecal LNP-DsiRNA-CCR2 interferes with cancer progression, we examined the femoral nerve, which connects the femur and the DRG. Because the anticancer effect of CCR2 DsiRNA is accompanied by decreased expression of pain-related neurotransmitters, reduced neuronal activity in the DRG and reduced pain-related behaviors, we quantified neurotransmitter content within the femoral nerve (Fig. 3A–C, Supplementary Fig. 2D and E). Histological analysis revealed increased levels of substance P (SP) and calcitonin gene-related peptide (CGRP) in the femoral nerve of cancer-bearing rats injected with the NC5 control relative to shams. CCR2 knockdown by repeated LNP-DsiRNA injection reduced SP and CGRP immunoreactivity in the femoral nerve innervating the bone tumor (Fig. 3B and C). CCL2 immunoreactivity, by contrast, was unchanged in cancer-bearing animals and was reduced below sham levels in LNP-DsiRNA-treated animals (Fig. 3A). Because SP and CGRP are released from peripheral nerve terminals^30^, we infer that these neurotransmitters are delivered into the tumor microenvironment.

**Figure 3.**
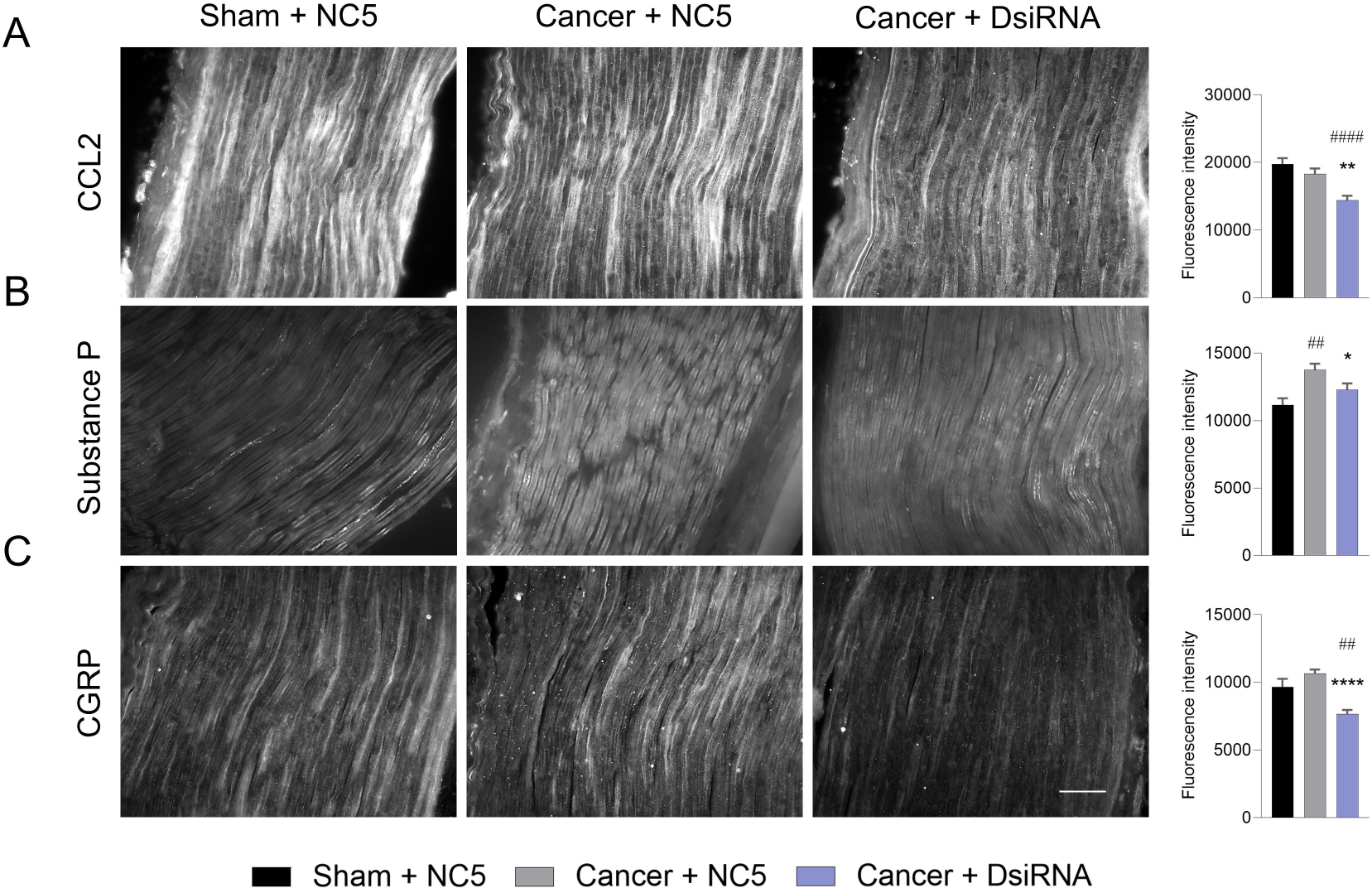
Substance P and CGRP levels are increased within the femoral nerve of cancer-bearing animals and are reduced following CCR2 silencing in the DRG. (A) CCL2, (B) substance P and (C) CGRP immunostaining in the ipsilateral femoral nerve of NC5-treated shams and of NC5- and CCR2 DsiRNA-treated cancer-implanted animals at POD 14 (n = 3 animals per group, 10 sections per animal). Data are presented as mean ± SEM. One-way ANOVA followed by Dunnett’s multiple comparisons test in (A–C). *P < 0.05, **P < 0.01, ***P < 0.001. # compared with the sham+NC5 group and * compared with the cancer+NC5 group.

### The reduced tumor burden induced by CCR2 silencing does not arise from altered tumor metabolism or cancer cell death

We next asked whether targeting CCR2 in the DRG decreases tumor burden by altering tumor metabolism. To this end, we performed positron emission tomography (PET) and measured fluorodeoxyglucose (^18^F-FDG) uptake in LNP-DsiRNA-treated tumor-bearing animals. *In vivo* PET revealed that LNP-DsiRNA did not reduce cancer-induced ^18^F-FDG uptake relative to NC5-injected animals, indicating comparable glucose uptake and therefore unaltered tumor metabolism on day 14 after implantation (Fig. 4A–D). ^18^F-FDG is not, however, specific for tumor metabolism and is also taken up by local inflammatory cells, which could mask a difference in tumor uptake.

**Figure 4.**
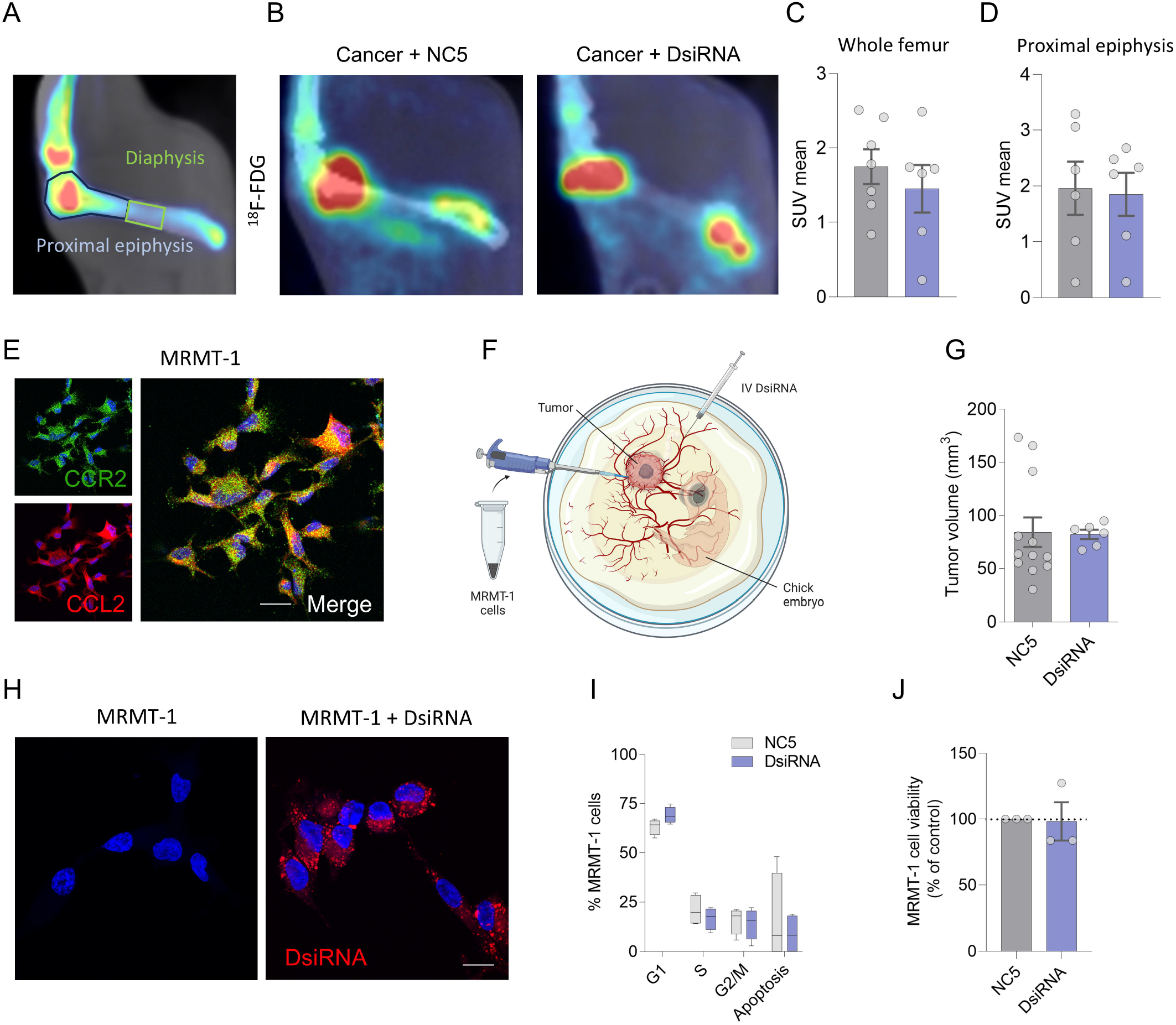
Tumor metabolism and cancer cell viability are not affected by CCR2 silencing. (A) Representation of the regions of interest analyzed on PET scans. (B) Representative images of ^18^F-FDG uptake in cancer+NC5 and cancer+DsiRNA animals. (C–D) Mean standardized uptake values (SUVmean) of ^18^F-FDG (cancer+NC5, n = 5; cancer+DsiRNA, n = 6). (E) Immunofluorescence analysis of CCR2 and CCL2 protein expression in MRMT-1 cells. Scale bar, 20 µm. (F) Schematic of the chorioallantoic membrane (CAM) model. (G) Tumor volume after NC5 and DsiRNA treatment in the CAM model. (H) MRMT-1 cell uptake of DID-labeled DsiRNA. Scale bar, 20 µm. (I) MRMT-1 cell cycle analysis by flow cytometry using the fluorescent DNA dye DAPI; effect of DsiRNA (n = 4; 250 ng) on the MRMT-1 cell cycle after 20 h of incubation compared with NC5 controls. (J) MRMT-1 cell viability assessed after 24 h of incubation with NC5 or DsiRNA (n = 5; 250 ng, performed in triplicate) using the MTT assay. Data are presented as mean ± SEM. Mann–Whitney test in (C, D, G and J); two-way ANOVA followed by Sidak’s multiple comparisons test in (I). *P < 0.05, **P < 0.01, ***P < 0.001; * compared with cancer+NC5.

To gain further insight into the mechanism driving the reduced tumor growth observed after CCR2 interference, we tested the effect of LNP-DsiRNAs on CCR2-expressing MRMT-1 cells in a chick embryo chorioallantoic membrane (CAM) model (Fig. 4E and F). In this model, cancer cells seeded onto a chick embryo form tumors whose response to intravenously delivered drugs can be assessed. Consistent with the *in vivo* PET data, CCR2 silencing (10 µg, 72 h) did not affect MRMT-1 tumor growth in the CAM model (Fig. 4G). We further tested the effect of CCR2 DsiRNA on MRMT-1 cells *in vitro*. LNP-DsiRNA uptake by MRMT-1 cells was successfully validated (Fig. 4H). Using DNA content-based flow cytometry, we monitored the progression of MRMT-1 cells through the cell cycle and found that LNP-DsiRNA did not alter the distribution of cells across cell cycle phases (Fig. 4I). Finally, an *in vitro* MTT assay revealed that anti-CCR2 DsiRNA did not significantly alter MRMT-1 cell survival (Fig. 4J). Taken together, these data show that the tumor-modulating effects of CCR2 silencing in the DRG *in vivo* are independent of any direct effect on cancer cells.

### Reduced tumor growth following CCR2 silencing is associated with reduced angiogenesis and bone degradation

Because of their high metabolic demand and growth rate, tumor cells rely on an increased nutrient supply delivered through enhanced blood perfusion. We therefore evaluated tumor neoangiogenesis using molecular imaging, exploiting the ability of RGD peptides to bind α^v^β^3^ integrins localized to newly formed blood vessels. We generated an α^v^β^3^-binding ^64^Cu-RGD peptide^31^ to monitor tumor neoangiogenesis by PET imaging (Supplementary Fig. 6A–E). The dynamic maximal standardized uptake value (SUVmax) of the tracer in the femur revealed significantly decreased ^64^Cu-RGD uptake in the tumor environment of LNP-DsiRNA-treated cancer-bearing animals relative to NC5-treated controls (Fig. 5A–C). This reduction in ^64^Cu-RGD retention within the tumor indicates reduced tumor angiogenesis in LNP-DsiRNA-treated rats.

**Figure 5.**
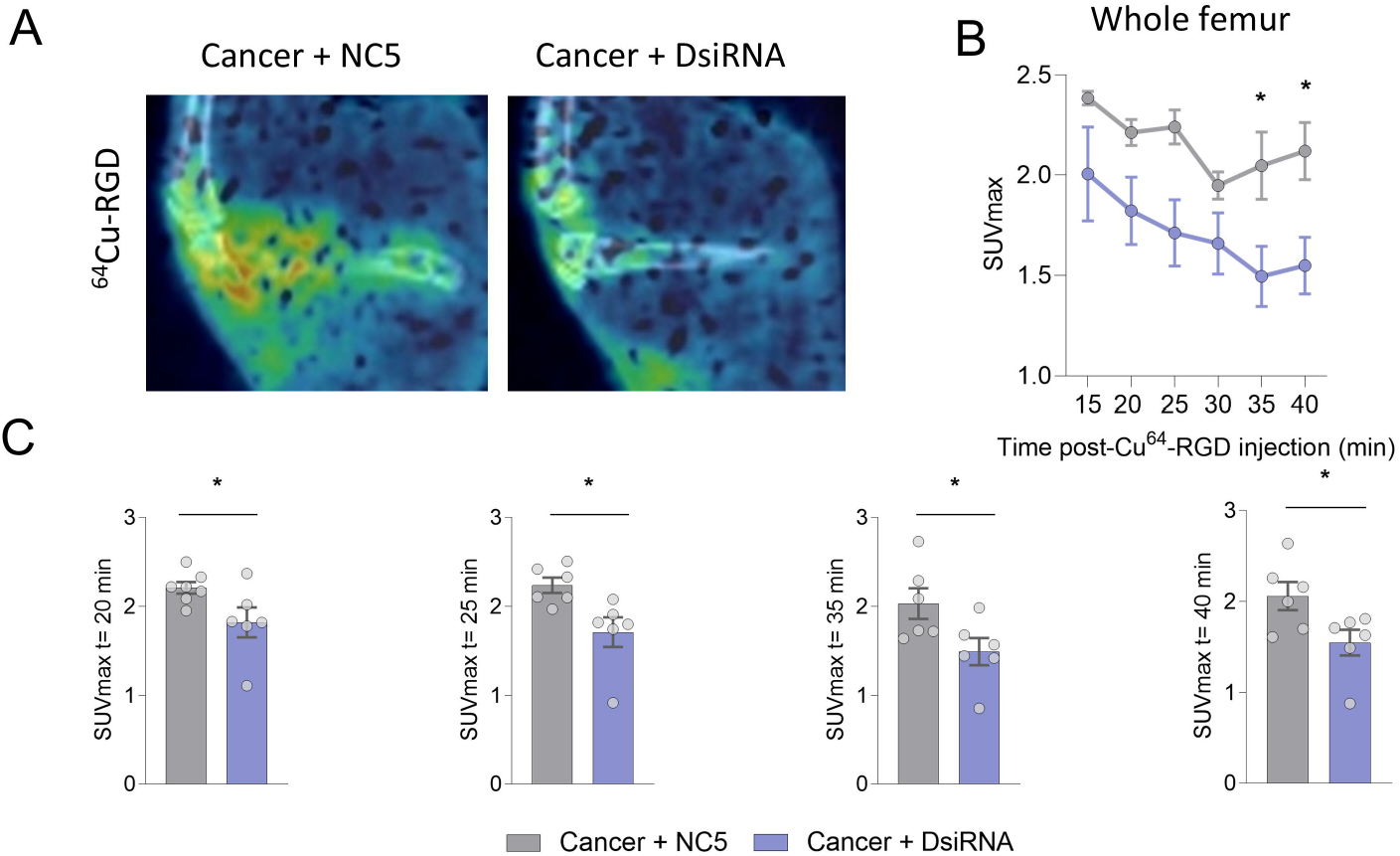
CCR2 silencing in the DRG reduces tumor angiogenesis. (A) Representative images of ^64^Cu-RGD uptake in cancer+NC5 and cancer+DsiRNA animals. (B–C) Maximal standardized uptake values of ^64^Cu-RGD PET scans (cancer+NC5, n = 5; cancer+DsiRNA, n = 6). Data are presented as mean ± SEM. Two-way ANOVA followed by Sidak’s multiple comparisons test in (B); unpaired t test in (C). *P < 0.05, **P < 0.01, ***P < 0.001; * compared with cancer+NC5.

Invasion of bone by tumor cells activates osteoclasts and accelerates bone remodeling, producing osteolytic lesions. The resulting increase in bone catabolism releases matrix-embedded growth factors and mediators that themselves display tumor-promoting activity^32^. We therefore asked whether the reduced tumor growth seen in LNP-DsiRNA-treated animals was accompanied by diminished bone remodeling. Three-dimensional reconstruction of micro-computed tomography (µCT) scans was used to analyze the morphological microstructure of femurs after implantation of MRMT-1 cells. Relative to non-implanted control femurs, cancer progression in NC5-treated animals caused severe bone remodeling: decreased bone volume, increased bone porosity, increased trabecular pattern factor and decreased bone volume/tissue volume (BV/TV) - histomorphometric parameters that together indicate a profoundly altered bone structure. Chronic silencing of CCR2 in the DRG returned bone remodeling towards sham levels (Fig. 6A–F).

**Figure 6.**
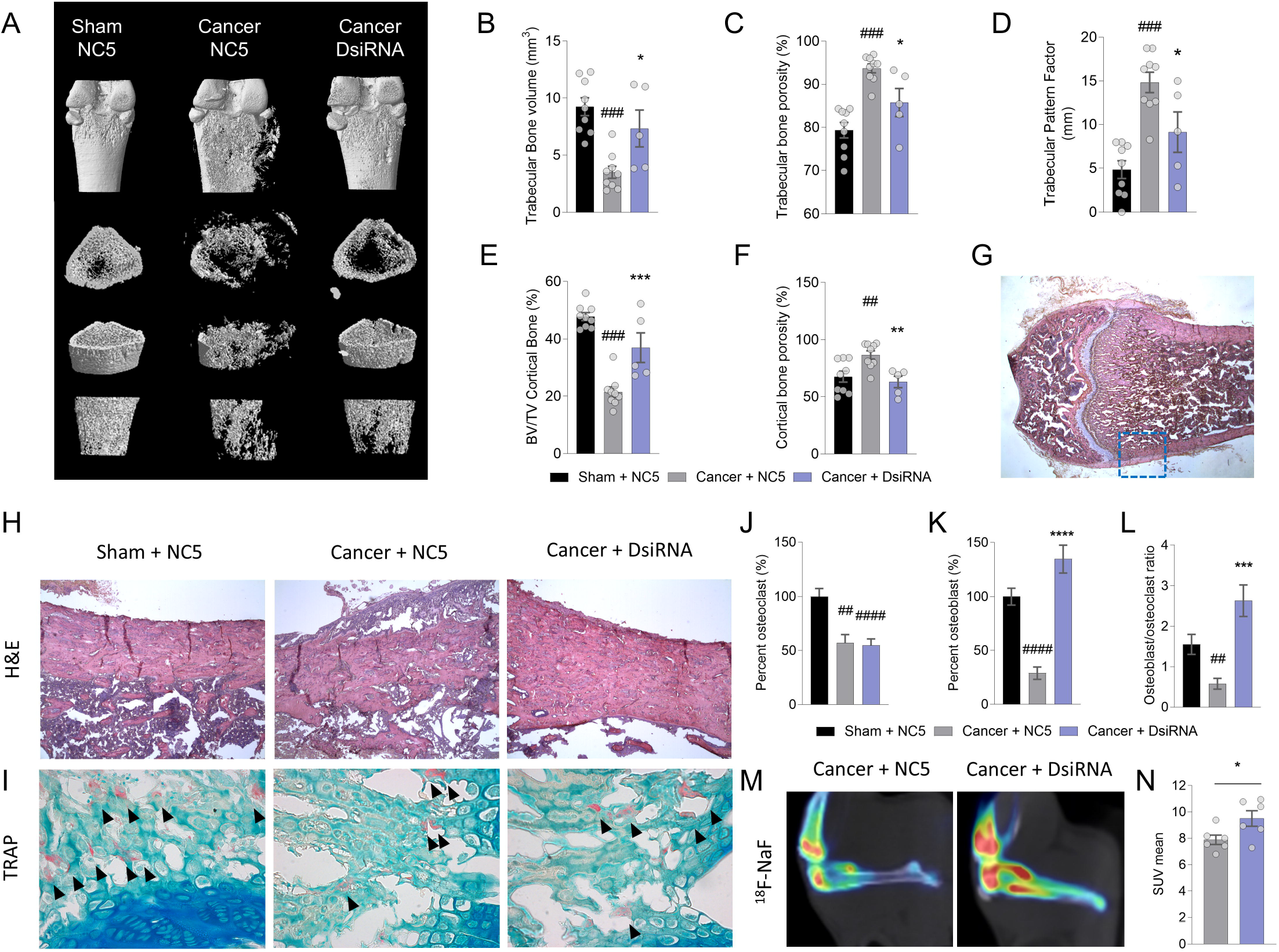
CCR2 silencing in the DRG prevents tumor-induced bone degradation. (A–F) Micro-computed tomography (µCT) analysis of the bone structure of sham-operated and tumor-implanted femurs (sham+NC5, n = 9; cancer+NC5, n = 9; cancer+DsiRNA, n = 5). (G–H) Representative histological images of hematoxylin and eosin (H&E)-stained sections showing cortical bone degradation that was prevented by LNP-DsiRNA treatment. (I) TRAP staining of bone sections. (J) Quantification of osteoclasts, (K) osteoblasts, and (L) the osteoblast-to-osteoclast ratio (sham + NC5, n = 4; cancer+NC5, n = 3; cancer+DsiRNA, n = 6; 10 sections analyzed per animal). (M) Mean standardized uptake values of 18F-sodium fluoride and (N) quantification (cancer+NC5, n = 5; cancer+DsiRNA, n = 6). BV/TV, bone volume/tissue volume. Data are presented as mean ± SEM. One-way ANOVA followed by Dunnett’s multiple comparisons test in (B–F); Kruskal–Wallis test followed by Dunn’s multiple comparison test in (I); unpaired t test in (J). *P < 0.05, **P < 0.01, ***P < 0.001. # compared with the sham+NC5 group and * compared with the cancer+NC5 group.

Histologically, hematoxylin and eosin (H&E) staining showed greater cortical bone porosity in NC5-treated cancer-bearing femurs than in sham-treated femurs. In contrast, chronic treatment with LNP-DsiRNA preserved cortical bone integrity, with reduced bone destruction within the tumor environment (Fig. 6G and H). At the cellular level, tartrate-resistant acid phosphatase (TRAP) staining revealed that the proportion of osteoclasts in the proximal epiphysis adjacent to the growth plate was reduced in NC5-treated cancer-bearing animals compared with NC5-treated sham animals and was not restored by LNP-DsiRNA treatment (Fig. 6I and J). LNP-DsiRNA treatment increased osteoblast numbers relative to LNP-NC5, resulting in a higher osteoblast-to-osteoclast ratio (Fig. 6K and L). Consistent with these findings, μCT and histological analyses showed preservation of bone architecture in LNP-DsiRNA-treated animals (Fig. 6A–H).

Bone metabolism was further assessed by PET imaging. Fluoride ions have high affinity for the bone matrix, and their deposition therefore serves as a marker of bone turnover. We used the radiotracer ^18^F-NaF to quantify bone turnover in NC5- and LNP-DsiRNA-treated rats on day 13 after cancer implantation. Increased ^18^F-NaF uptake following CCR2 silencing in the DRG (Fig. 6M and N) indicates increased bone formation, consistent with the preserved bone architecture observed by µCT. Collectively, these results establish that CCR2 silencing in the DRG limits tumor angiogenesis and maintains bone integrity.

### Substance P and CGRP promote expansion of the macrophage population in the tumor niche

We next sought to determine how CCR2 silencing in the DRG could affect tumor angiogenesis and tumor-induced bone remodeling in the femur. As shown in Figure 3, substance P and CGRP levels were increased in the femoral nerve arising from the lumbar DRG of cancer-bearing animals, and both were normalized by CCR2 DsiRNA. The peripheral release of these neurotransmitters and their influence on local immunity are widely documented in the skin during fungal and other infections^33^, a process referred to as neurogenic inflammation. We therefore examined the direct effect of substance P and CGRP on the cellular components of the tumor niche using an *in vitro* WST-1 assay (Fig. 7A–C). Neither neurotransmitter affected the MRMT-1 tumor cells used in our *in vivo* bone cancer model. Consistent with this, MRMT-1 cells expressed calcitonin receptor-like receptor (CRLR) but not receptor activity-modifying protein 1 (RAMP1), which is required for CGRP signaling, and did not express the substance P receptor neurokinin 1 (NK1R) (Supplementary Fig. 7A–C). Endothelial cell and osteoblast proliferation were likewise unaffected by substance P and CGRP, whereas both neurotransmitters exerted a significant pro-proliferative effect on monocytes/macrophages (Fig. 7A–C). To test whether this translates to the *in vivo* setting, we analyzed tumor-infiltrating macrophages by flow cytometry following intrathecal LNP-DsiRNA injection. Consistent with the *in vitro* data, macrophage numbers were significantly reduced by CCR2 silencing in the DRG (Fig. 7D). Disrupting CCR2 in the DRG therefore reduces neurotransmitter transport along the femoral nerve innervating the tumor and lowers the number of macrophages that are likely to promote tumor growth, tumor angiogenesis and tumor-induced bone resorption.

**Figure 7.**
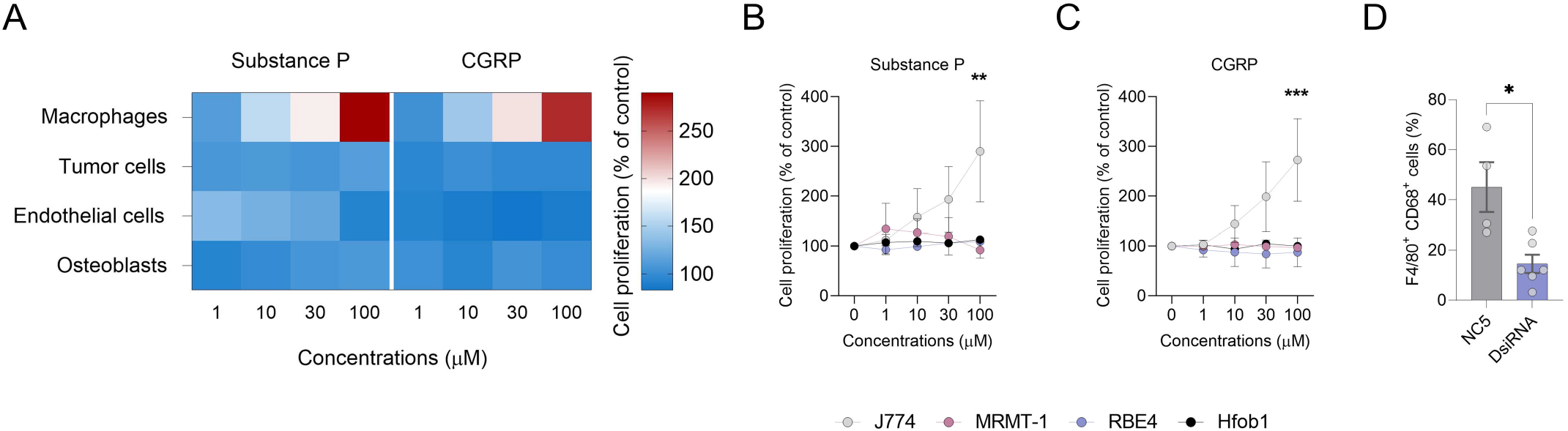
Substance P and CGRP induce macrophage expansion, and targeting CCR2 in the nervous system reduces the number of TAMs. (A–C) Dose–response of macrophage (J774), tumor cell (MRMT-1), endothelial cell (RBE4) and osteoblast (hFOB1.19) proliferation assessed after 24 h of incubation with substance P or CGRP (n = 3; performed in triplicate) using the WST-1 assay. (D) Flow cytometric analysis of F4/80^+^ CD68^+^ tumor-associated macrophage numbers in NC5- and DsiRNA-treated animals. Data are presented as mean ± SEM. Two-way ANOVA followed by Sidak’s multiple comparisons test in (B and C); Mann–Whitney test in (D). *P < 0.05, **P < 0.01, ***P < 0.001.

## Discussion

The bone niche offers a rich and unique environment for metastatic seeding and tumor growth. Within this microenvironment, multiple components contribute to tumor expansion, including tumor-infiltrating nerves, bone-degrading osteoclasts, endothelial cells and infiltrating immune cells. We report that tumor progression within the femur activates DRG neurons, leading to antidromic transport of substance P and CGRP along the tumor-innervating femoral nerve. Silencing CCR2 in the DRG decreased neuronal activation and antidromic neurotransmitter transport, reduced macrophage numbers and blunted tumor angiogenesis and osteolysis, resulting in a reduced tumor burden. The critical involvement of neurogenic inflammation in tumor promotion was further supported by femoral nerve axotomy, which hindered tumor growth, and by intrathecal injection of exogenous CCL2, which promoted tumor progression.

Overwhelming evidence points to a role for the autonomic nervous system in tumorigenesis^5,8,14^. The nervous system modulates tumor initiation and development in a manner that depends on the type of innervation and the neurotransmitters involved: the sympathetic and enteric nervous systems promote cancer progression, whereas the parasympathetic system hinders tumor growth^4,13,34,35,36,37^. The role of sensory innervation in tumor promotion has also been investigated, with reports that capsaicin-induced denervation delays pancreatic cancer progression and that surgical denervation of touch domes limits skin cancer development^6,38,39^, and with CGRP playing a substantial role in tumor progression by shaping immunosurveillance^40^.

Here, we propose that neurotransmitters transported antidromically as a consequence of afferent neuron activation by bone tumors drive tumor progression. The femur is densely innervated by peptidergic nerve endings, and substance P and CGRP contribute to angiogenesis and to bone metabolic homeostasis^41,42,43^. Antagonists of NK1, the high-affinity receptor for substance P, block tumor angiogenesis and tumor growth^44^. Similarly, CGRP displays proangiogenic properties in tumors, and CGRP antagonism with the FDA-approved antimigraine drug rimegepant enhances the efficacy of antitumor therapy^40,43^. In our study, CCR2 silencing in DRG neurons reduced pain as well as substance P and CGRP transport into the femoral nerve.

Recent work has further highlighted the role of sensory neuropeptides in tumor progression. Neuronal substance P was shown to promote breast tumor growth, invasion and metastasis through tumoral TACR1 and an extracellular single-stranded RNA–TLR7 axis, with the TACR1 antagonist aprepitant suppressing growth and metastasis in multiple models^62^. Similarly, nociceptor-derived CGRP promotes gastric tumor progression through RAMP1 signaling^63^ and drives pancreatic ductal adenocarcinoma progression by acting on cancer-associated fibroblasts to suppress natural killer cells^64^. CGRP also remodels tumor-draining lymph nodes into an immunosuppressive microenvironment characterized by M2-like polarization of tumor-associated macrophages^65^, mirroring the macrophage-dependent mechanism we describe here. In bone, periosteal CGRP-positive sensory fibers are enriched in mice bearing bone metastases, serum CGRP is elevated in patients with bone metastases, and blockade of sensory neuron-derived CGRP reduces bone metastatic progression^66^. Notably, MRMT-1 cells express CRLR but lack RAMP1, and neither CGRP nor substance P altered their proliferation, indicating that in our model these neuropeptides act on the stroma rather than on the tumor cells themselves. This distinction from models in which CGRP is directly mitogenic^66^ likely reflects tumor cell-intrinsic differences in RAMP1 and TACR1 expression and argues that the target of neuropeptide signaling should be established for each tumor model^67^.

Acting through its cognate high-affinity receptor CCR2, the chemokine CCL2 exerts pleiotropic effects, including immune cell recruitment and the induction of peripheral and nervous system inflammation^46,47^. CCL2 is an important and well-characterized neuromodulator in both the peripheral nervous system and the spinal cord. CCL2 released from primary afferent neurons increases nociceptor sensitization, synaptic transmission in the dorsal horn and neuroinflammation leading to nociception^48,49,50,51^. Antagonists acting at CCR2 alleviate neuropathic and inflammatory pain in rodent models^26,52,53,54,55^, and allosteric CCR2-targeting pepducins relieve bone cancer pain without promoting tumor progression^16^. Here, we explored the consequences of sustained neuroinflammation and peripheral neurogenic inflammation for tumor progression, and identify CCR2 silencing in sensory neurons as a therapeutic strategy.

In this study, we provide evidence that targeting neuroimmune communication by silencing CCR2 signaling in the DRG limits tumor growth. The complex interplay between metastatic bone tumors and immune cells is a detrimental interaction that drives bone tumor growth, pathological osteolysis and pain, and the convergence of independent approaches now targeting this tumor– nerve crosstalk in bone^71^ supports the neuroimmune axis as a promising treatment strategy.

## Materials and Methods

### Animals

Female Sprague–Dawley rats (150–175 g; Charles River Laboratories) were maintained on a 12 h light/dark cycle with access to food and water *ad libitum*. All animal procedures were approved by the ethical animal care committee of the Université de Sherbrooke, in compliance with the policies and directives of the Canadian Council on Animal Care and with the ARRIVE guidelines.

### Axotomy of the femoral nerve

Unilateral denervation of the femoral nerve was performed under isoflurane anesthesia. The skin was shaved and disinfected by alternating three washes with 70% ethanol and three washes with the detergent dexidine 4 (Atlas Laboratories, #LAT918140). A minimal (1 cm) opening in the abdominal cavity was made midway between the diaphragm and the pubis by cutting the skin and the superficial abdominal muscles. The psoas major and psoas minor were separated by gently tearing the fasciae. The femoral nerve was exposed and a 5 mm segment was excised. The abdominal muscles were sutured with Monocryl suture thread (4-0; Ethicon #Y494G) and the dermis with Prolene suture thread (5-0; Ethicon #8696G). The skin was washed with 3% hydrogen peroxide, and animals received subcutaneous injections of sterile ampicillin (100 mg/kg) twice daily for three days and of meloxicam (5 mg/kg) once daily for three days. Sham-operated rats underwent the same procedure without transection of the femoral nerve. Femoral nerve axotomy did not produce a detectable change in motor function or limb use, and nerve transection was confirmed at dissection.

### Cell culture

MRMT-1 rat breast carcinoma cells were kindly provided by the Cell Resource Center for Biomedical Research, Institute of Development, Aging and Cancer (Tohoku University), and were maintained in RPMI 1640 medium supplemented with 10% FBS and 2% penicillin/streptomycin at 37 °C in a humidified atmosphere containing 5% CO^2^.

### Cancer implantation surgery

Rats were randomly assigned to the cancer group or the sham surgery group. Syngeneic MRMT-1 breast cancer cells were implanted surgically as described by Doré-Savard et al.^56^. Briefly, 30,000 cells diluted in 20 µL Hank’s balanced salt solution (HBSS) were injected into the medullary cavity of the femur of female rats after a minimal opening had been made with a microdrill. The hole was then sealed with dental amalgam. No postoperative analgesia was administered, to avoid interference with pain assessment. Sham-operated rats underwent all surgical procedures, but no cancer cells were delivered into the femoral bone marrow.

### CCL2-driven neuroinflammation

Animals were randomly assigned to experimental groups and injected intrathecally with saline or CCL2 (1 µg/rat, dissolved in 0.1% BSA in sterile saline; R&D Systems #3144-JE-025/CF) on postoperative days (PODs) 3, 7 and 12 under light isoflurane anesthesia.

### Measurement of tumor growth in vivo

Tumor size was monitored daily by palpation of the femur according to the following criteria: 0 = no tumor; 1 = slight distention of the femoral epiphysis; 2 = tumor mass around the epiphysis; 3 = tumor mass bulging 3 mm out of the epiphysis; 4 = tumor mass protruding 5 mm out of the epiphysis; and 5 = tumor mass bulging out of the epiphysis and diaphysis. Representative images are shown in Supplementary Fig. 1A. A higher score indicates a larger tumor.

### Immunostaining

After perfusion, the L1-L3 spinal cord, ipsilateral L1 to L3 DRG and femoral nerves were collected, post-fixed in 4% paraformaldehyde at 4 °C for 24 h and cryoprotected in 30% sucrose in PBS at 4 °C for 48 h. Frozen tissues were embedded at −35 °C in O.C.T. compound; 30 µm transverse spinal cord sections were cut on a Leica SM220R sliding microtome and 20 µm DRG and nerve sections on a cryostat, and collected on gelatin-coated SuperFrost Plus slides. Sections were blocked (10% NGS, 1% BSA, 0.05% Tween-20, 0.1% Triton X-100 in PBS) and incubated in 0.3 M glycine containing 0.2% Tween-20. Sections were labeled with a chicken anti-CCR2 antibody (1:200; Aves Labs, custom), chicken anti-CGRP (1:500, Millipore #AB5705), guinea-pig anti-substance P (1:500, Neuromics #GP14110) in blocking buffer for 24 h at room temperature, then incubated with a fluorophore-conjugated secondary antibody (1:500, AlexaFluor 488, 568, 647; Invitrogen). All slides were coverslipped with ProLong Diamond Mountant. Fluorescence images of femoral nerve slices were acquired at 20x using a Leica DM4000 microscope equipped with a Leica DFC350FX camera using the same acquisition parameters. Fluorescence intensity quantifications were performed using ImageJ software. 9-10 sections were analyzed per animal, for a total of three animals per treatment condition. Representative fluorescence images were acquired either on an Olympus FV1000 confocal microscope or on a Leica DM4000. The anti-CCR2 antibody was validated in CCR2-positive (spleen) and CCR2-negative (heart) organs, as well as in CCR2-positive and CCR2-negative cells^16^.

### DsiRNA design

All chemically modified DsiRNAs described in this study were synthesized by Integrated DNA Technologies Inc. (IDT). The identity of each duplex was verified by electrospray ionization mass spectrometry (ESI-MS) and was within ± 0.02% of the predicted mass. Purification was performed by high-performance liquid chromatography, and products were > 90% pure. Duplexes were prepared as sodium salts, with different recognition sites within the rat CCR2 (rCCR2) mRNA and different chemical modifications.

### LNP-DsiRNA formulation

LNP-DsiRNAs were formulated using a proprietary lipid mixture containing an ionizable cationic lipid, supplied as Neuro9™, and DsiRNAs were encapsulated using a microfluidic system for controlled mixing on a NanoAssemblr™ instrument (Precision Nanosystems)^57^. The quality and uniformity of the lipid nanoparticles were assessed with a particle size analyzer (Malvern Zetasizer). The Z-average size for each batch used was 60–70 nm, with a polydispersity index of 0.03–0.08. Encapsulation efficiency was determined with a RiboGreen assay (Invitrogen) and was greater than 90%.

### Compound administration

After cancer cell implantation, animals were randomly assigned to the control (vehicle, NC5) or treatment (DsiRNA) group. Rats received daily administration of either DsiRNA (297 pmol/rat in PBS, intrathecally) or the NC5 control from day 11 to day 14 under light anesthesia. Injections were performed two hours before behavioral testing.

### Mechanical allodynia

Rats were acclimatized to the von Frey apparatus for three consecutive days. The mid-plantar surface of the ipsilateral and contralateral hind paws was stimulated with von Frey hairs of logarithmically increasing stiffness by an experimenter blinded to treatment. Hairs were held for 7 s with intervals of several seconds between stimulations. Stimuli were presented consecutively, descending after a behavioral response and ascending otherwise. Testing ended after a negative response to the stiffest hair (15 g) or after four stimulations following the first positive response. The 50% paw withdrawal threshold was interpolated using the formula 50% g threshold = 10^[Xf + kδ], where Xf is the value (in log units) of the final von Frey hair used, k is the tabular value for the pattern of positive/negative responses, and δ is the mean difference (in log units) between stimuli.

### Paw retroflexion scores

Ipsilateral paw retroflexion was scored as follows: 0 = full contact with the floor; 1 = light contact with the floor; 2 = light contact with the floor due to paw deformation; 3 = curved paw with only the tips of the toes touching the floor; and 4 = curved paw with no contact with the floor. A higher score indicates more pain.

### Dynamic weight bearing

The dynamic weight bearing (DWB) device (Bioseb) consists of a Plexiglas enclosure (22 × 22 × 30 cm) with a floor sensor composed of 44 × 44 captors (10.89 mm^2^ per captor). As previously described^58^, the animal moved freely within the apparatus for 5 min while pressure data and live recordings were transmitted to a laptop computer via a USB interface. Raw pressure and visual data were collated with DWB software v1.4.2.98.

### In vivo PET and CT imaging

Positron emission tomography (PET) imaging was performed on a Triumph™ PET/CT dual-modality platform (Gamma Medica, Inc.) equipped with an avalanche photodiode-based digital PET scanner with a 7.5 cm axial field of view. The scanner achieved a transaxial spatial resolution of 1.2 mm and a detection efficiency of 2.6% with an energy window of 250–650 keV.

Rats received approximately 20 MBq of ^18^F-sodium fluoride (^18^F-NaF) and approximately 20 MBq of ^18^F-fluorodeoxyglucose (^18^F-FDG) by intravenous injection (200 μL at 500 μL/min) on PODs 13 and 14, respectively. Thirty minutes after radiotracer administration, the hind knee joints were aligned at the radial and axial centers of the scanner field of view using a custom-built paw support designed to position both limbs stably and reproducibly, and radiotracer accumulation in the target tissues was monitored by 30 min of static imaging. Six hours after ^18^F-FDG injection, a 30 s blank scan was performed to assess residual FDG signal, followed by injection of 35 MBq of ^64^Cu-RGD; tracer accumulation in the femur was then monitored by 45 min of dynamic imaging.

After each scan sequence, a cylindrical phantom containing a known quantity of free radiotracer, matched to the approximate size of the rat (∼30 mL), was used to obtain a calibration factor converting counts per second into absolute activity in mBq, from which SUVmean and SUVmax values were derived. A density of 1 g/cc was used to convert fractional uptake per volume into SUVmean and SUVmax values. Images were reconstructed with the Triumph PET/CT software using a 3D-MLEM algorithm with 20 iterations, a span of 63 and a field of view of 80 mm, giving a final matrix resolution of 160 × 160 × 128 and a voxel size of 0.5 × 0.5 × 0.597 mm^3^.

CT acquisition was performed over 360° using 512 projections, a time frame of approximately 250 ms per projection, a field of view of 84.57 mm, a source peak voltage of 60 kVp at 230 μA and 2 × 2 pixel binning. The single frame was reconstructed into 0.165 × 0.165 × 0.165 mm^3^ voxels. CT image reconstruction was performed as previously described^59^.

### PET and CT image visualization and analysis

PET images were coregistered to the animal’s CT image using rigid transformation with the cross-modality 3D image fusion tool implemented in PMOD 3.8 (PMOD Technologies Ltd.). Regions of interest (ROIs) were delineated on CT images of the cancer-implanted limb or of the contralateral limb as controls. ROIs were drawn on the femur to define the diaphysis and the proximal and distal epiphyses relative to the implantation site; the extraosseous tumor mass was also included. ROIs were drawn on consecutive coregistered PET images. Raw data were extracted from these ROIs, and the mean and maximal standardized uptake values (SUVmean and SUVmax) were calculated as follows:

SUVmean = mean uptake value / (injected dose [MBq] × animal weight [kg]) SUVmax = maximum uptake value / (injected dose [MBq] × animal weight [kg])

SUV values were further corrected for injection time and radioactive decay according to: Corrected SUV = SUV × EXP−λ(τPET − τInjection), where λ = Ln(2)/τ1/2

### Chorioallantoic membrane (CAM) assays

Fertilized White Leghorn chicken eggs were obtained from the Public Health Agency of Canada (Ottawa, ON, Canada). The Ethics Committee on Animal Research at the Université de Sherbrooke approved the protocols, and all embryo-related procedures followed the regulations of the Canadian Council on Animal Care. CAM assays were carried out essentially as described^60^, with minor modifications. On embryonic day (ED) 9, a suspension of MRMT-1 cells (1 × 10^6^ cells) was combined with standard Matrigel basement membrane matrix (VWR, #CACB354234) at a 1:1 ratio in a total volume of 20 µL. The number of cells implanted was chosen on the basis of preliminary dose–response experiments. On ED 13, CCR2-6 S3AS12 or NC5 control DsiRNA (10 µg) was injected intravenously into the CAM. At ED 16, chick embryos were sacrificed by decapitation. Tumors were removed and imaged, and tumor volumes were calculated as Dd^2^/2, where D is the largest diameter and d is the smallest diameter.

### ApoE-dependent penetration of lipid nanoparticles

MRMT-1 cells were treated with DsiRNA-containing lipid nanoparticles (5 µg/ml) labeled with DID dye, in the presence or absence of recombinant ApoE4 (10 µg/ml; PeproTech #350-04) for 24 h. Cells were fixed with 4% PFA for 30 min, rinsed and incubated in 0.1% glycine in PBS for 20 min. Cells were then incubated with 0.625 µg/ml DAPI, rinsed and coverslipped with ProLong Gold mountant (Invitrogen #P36930).

### Analysis of the cell cycle by flow cytometry

MRMT-1 cells (30,000) were seeded in triplicate in 48-well plates in RPMI 1640 medium supplemented with 10% FBS and 2% penicillin/streptomycin. Twelve hours after seeding, cells were serum-starved for 24 h for synchronization. Fresh medium containing 2% FBS supplemented with NC5 or DsiRNA was then added. For NC5 and DsiRNA uptake, 10 µg/ml ApoE (PeproTech #350-04) was added. After 20 h of incubation, cells were detached with Accutase, which was quenched with medium containing 2% FBS. Cells were pelleted in a microcentrifuge and resuspended in 50 µL FACS buffer (PBS containing 2% FBS). Cells were fixed with 70% ethanol at −20 °C, added dropwise with gentle vortexing, and incubated on ice for 30 min. Cells were then pelleted, resuspended in 150 µL FACS buffer containing 2 µg/mL DAPI and incubated on ice for 30 min. DAPI fluorescence was analyzed on a Beckman Coulter CytoFLEX 30 flow cytometer at a flow rate of 30 µL/min using the blue channel. The percentage of cells in each phase of the cell cycle was calculated with ModFit software.

### MTT assay

MRMT-1 cells (15,000) were seeded in triplicate in 96-well plates in phenol red-free RPMI 1640 medium supplemented with 10% FBS and 2% penicillin/streptomycin. Twelve hours after seeding, 100 µl of fresh serum- and phenol red-free medium containing NC5 or DsiRNA was added. For NC5 and DsiRNA uptake, 10 µg/ml ApoE (PeproTech #350-04) was added. After 72 h of incubation, 10 µl of MTT reagent (5 mg/ml) was added to each well and plates were incubated for 4 h at 37 °C and 5% CO^2^. Cells were centrifuged at 1,000 × g for 5 min and the medium was carefully removed. DMSO (100 µl) was added to dissolve the formazan crystals, and the optical density (OD) of each well was measured at 450 nm on a GENios Pro plate reader (Tecan). Cell viability was calculated as (OD450 nm treatment / OD450 nm scrambled control) × 100.

### Synthesis of the RGD-NOTA-64Cu radionuclide

Unless otherwise stated, all reactions were performed under a nitrogen atmosphere. All solvents (ChemImpex International), amino acid derivatives, coupling reagents (ChemImpex International), piperidine and N-methylpyrrolidinone (A&C American Chemicals Ltd.) were HPLC grade. All other reagents were purchased from Sigma-Aldrich. Water-sensitive reactions were performed in anhydrous solvents. Peptide synthesis was performed manually using Fmoc-Gly-2-Cl-Trt chloride resin (0.80 mmol g^-1^, Matrix Innovation). Two grams of resin was treated with a solution of Fmoc-Gly-OH (0.6 mmol) and diisopropylethylamine (1.2 mmol) in 30 mL dichloromethane for 1.5 h. Two milliliters of a 1:1 mixture of MeOH and diisopropylethylamine was added and the mixture was agitated for 5 min. The resin was washed five times each with dichloromethane and isopropyl alcohol, then dried overnight. Using 1.0 g of this resin (0.3 mmol) with a 5-fold excess of amino acid, 1-[bis(dimethylamino)methylene]-1H-1,2,3-triazolo[4,5- b]pyridinium 3-oxide hexafluorophosphate (HATU) and a 10-fold excess of diisopropylethylamine in 12 mL N,N-dimethylformamide (DMF), Fmoc-Arg(Pbf)-OH was coupled. The resin was washed five times with DMF and the Fmoc protecting group was removed with 20% piperidine in DMF for 15 min. After five further washes with DMF, Dde-Lys(Fmoc)- OH and NOTA bis t-butyl ester (Chematech) were coupled using the same coupling/deprotection sequence. The Dde group on the alpha amine of the lysine was removed by treatment with 2% hydrazine in DMF (3 × 2 min) and the resin was washed seven times with DMF. Fmoc-D-Tyr-OH and Fmoc-Asp(OtBu)-OH were then coupled as described above, the final Fmoc group was removed, and the resin was washed three times with DMF and five times with dichloromethane. After cleavage with 30% hexafluoropropan-2-ol in dichloromethane for 2.5 h, filtration and solvent removal under reduced pressure on a rotary evaporator, a crude, partially protected peptide was obtained. This crude peptide was dissolved in 200 mL tetrahydrofuran and macrocyclized with DEPBT (215 mg, 1.5 eq) and diisopropylethylamine (250 µL, 3.0 eq) overnight. The tetrahydrofuran was removed on a rotary evaporator and the crude macrocycle was dissolved in 20 mL dimethylformamide. The protected macrocycle was obtained after elution on a carbonate cartridge (Silicycle) followed by evaporation on a speed vac system. The macrocycle was dissolved in 20 mL TFA/water (95/5) for 2.5 h to remove the protecting groups, and the TFA was evaporated under reduced pressure. The product was purified on a mass-triggered preparative system (Waters) equipped with an X-Select CSH C18 column (5 µm, 10 × 100 mm). Pure fractions were lyophilized to yield the final product as a white powder. The peptide was analyzed on an Acquity UPLC-MS system class H (Acquity UPLC protein BEH C18 column, 2.1 mm × 50 mm, 1.7 μm particles with 300 Å pores) coupled to an SQD2 mass spectrometer (Waters) and a PDA UV–visible detector, using the following gradient at a flow rate of 800 µL/min: 0 min, 5% ACN; 0.2 min, 5% ACN; 1.5 min, 95% ACN; 1.8 min, 95% ACN; 2.0 min, 5% ACN; 2.5 min, 5% ACN.

Our cyclotron facility provides ^64^Cu isotopes for research purposes on a routine basis using a target system developed in collaboration with ACSI (Richmond, BC, Canada). ^64^Cu was prepared on a TR-19 or TR-24 cyclotron (ACSI) via the ^64^Ni(p,n)^64^Cu reaction using an enriched ^64^Ni target electroplated on a rhodium disc. ^64^CuCl^2^ was recovered from the target and converted to ^64^Cu- acetate by dissolution in ammonium acetate (0.1 M, pH 5.5) followed by evaporation to dryness. For labeling studies with ^64^Cu(OAc)^2^, the peptide (3–5 nmol) was dissolved in 1 M ammonium acetate buffer (pH 7.4), the radiometal (150–250 MBq) was added, and the mixture was incubated for 40 min at room temperature. Radiolabeling was monitored by UPLC. The labeled product was purified on a C-18 Sep-Pak (Waters Corporation) by elution first with HPLC water and then with acetonitrile. The peptide fraction in acetonitrile was collected and evaporated, and the ^64^Cu peptide was counted in a Capintec radioisotope calibrator to calculate the specific activity of the product, estimated at 1800–2000 Cu/mmol.

### Ex vivo µCT

Anesthetized rats were perfused intra-aortically with 100 ml saline followed by 500 ml 4% paraformaldehyde. Ipsilateral femurs were removed, post-fixed for 48 h in the same solution and washed in PBS for *ex vivo* µCT. Scans were performed at the McGill Bone Center on a high-resolution desktop micro-CT scanner as previously described^61^. Rat femurs were scanned at an X-ray source power of 45 keV/222 µA and a resolution of 11.25 µm/pixel. µCT images were reconstructed with NRecon (v1.6.1.3), and 3D analyses were performed with CT-Analyzer (v1.10.0.2), both provided by SkyScan. The volume of interest (VOI) for cortical plus trabecular bone was defined as the total (tissue) volume, including cortical bone, trabecular bone and any intervening spaces, over a range of 5.626 mm (201 cross-sections) starting from the growth plate in the distal femur. The trabecular VOI was defined as the total (tissue) volume, including all trabecular bone and any spaces, within the same 5.626 mm range starting from the growth plate in the distal femur.

### Bone histology

After µCT scanning, femurs were embedded in a 50% methyl methacrylate (MMA)/50% glycol methacrylate (GMA) mixture and cut into 6 μm sections. MMA-GMA was removed by two 30 min immersions in 2-methoxyethyl acetate followed by two 5 min xylene baths, and samples were rehydrated through graded ethanol. Samples were then equilibrated in PBS for 15 min. Hematoxylin and eosin (H&E) staining was performed to count cuboidal bone-lining osteoblasts. Osteoclasts and other mononuclear TRAP-positive cells were stained with Naphthol AS-TR phosphate, sodium nitrite, sodium tartrate and pararosaniline hydrochloride in acetate buffer (pH 5.0). Images were acquired on a Leica DM4000 microscope equipped with a Leica DFC350FX camera using identical acquisition parameters across groups. Eight to ten sections were analyzed per animal (n = 4 sham+NC5, n = 3 cancer+NC5, n = 6 cancer+DsiRNA).

### WST-1 assay

MRMT-1, J774 macrophage, RBE4 endothelial, and hFOB1.19 osteoblast cells (15,000 cells/well) were seeded in triplicate in 96-well plates in their respective growth media: RPMI 1640 (MRMT- 1), DMEM (J774), DMEM supplemented with 1 ng/mL basic fibroblast growth factor (RBE4), or a 1:1 mixture of DMEM and Ham’s F-12 (hFOB1.19), each containing 10% fetal bovine serum (FBS) and 2% penicillin/streptomycin. All cell lines were treated in parallel. Twelve hours after seeding, 100 µl of fresh medium containing 2% FBS supplemented with 1, 10, 30 or 100 µM CGRP (Bachem #50-259-898), substance P (Sigma #S6883) or the corresponding vehicle (PBS) was added. After 24 h of incubation, 10 µl of WST-1 reagent (5 mg/ml) was added to each well and plates were incubated for 4 h at 37 °C and 5% CO^2^. The optical density (OD) of the formazan dye generated by conversion of the WST-1 tetrazolium salt by mitochondrial enzymes was measured at 450 nm for each well on a GENios Pro plate reader (Tecan). Cell viability was calculated as (OD450 nm treatment / OD450 nm scrambled control) × 100.

### Isolation of tumor-infiltrating cells

Anesthetized rats were decapitated and the ipsilateral femur was collected and crushed with a mortar and pestle. Tumor tissue was finely minced and digested for 25 min with 0.025% collagenase A (#10103586001; Roche) and 1 mM EDTA in PBS at 37 °C with frequent agitation and trituration. Cells were filtered through a 70 µm cell strainer (Fisherbrand #22363548) and washed with RPMI 1640 medium supplemented with 10% FBS. Cells were centrifuged at 450 × g for 6 min at 4 °C, resuspended and incubated in erythrocyte lysis buffer at room temperature for 2 min. Lysis was stopped with cold RPMI medium, after which cells were filtered through a 70 µm cell strainer and centrifuged at 450 × g for 6 min at 4 °C. The cell pellet was resuspended in FACS buffer (10% NGS, previously decomplemented at 56 °C for 30 min, plus 1 mM EDTA in PBS).

### Flow cytometry assays

Dissociated cells were briefly fixed with 2% paraformaldehyde for 30 min at 4 °C, centrifuged at 1,600 rpm at 4 °C and washed with cold FACS staining buffer. Cells were centrifuged at 1,600 rpm at 4 °C, resuspended in 100 μl FACS buffer and incubated with an anti-CD32 antibody (1:20; BD Biosciences #550271) for 30 min at 4 °C to block non-specific Fc receptor binding. Multicolor immunostaining was performed with F4/80-AlexaFluor 647 (1:40; Bioss Antibodies #BS-7058R- A647), CD163-FITC (1:40; Biorbyt #orb434303), CD68-PE-Cy7 (1:40; Abcore #AC12-0231-17), CD3-APC-Cy5.5 (1:40; Abcore #AC12-0212-04), CD4-PerCP-eFluor710 (1:40; eBioscience #46- 0040-82) and CD8a-PE (1:40; BioLegend #200607) antibodies in FACS buffer containing 2 µg DAPI for 1 h at 4 °C. Cells were rinsed, resuspended in FACS buffer containing DAPI and analyzed on a Beckman Coulter CytoFLEX flow cytometer (Beckman Coulter, Brea, CA, USA). The specificity of each antibody was validated on activated rat splenocytes and on peritoneal macrophages (2% thioglycollate intraperitoneally, 3 mL, 72 h) isolated as described in the supplementary material. Single-stained compensation beads (VersaComp Antibody Capture Beads #B22804) were used to measure the spillover of each dye into neighboring channels for multicolor compensation, and an electronic compensation matrix was applied to correct this crosstalk. Positive and negative selection gates were set using fluorescence-minus-one and unstained controls. Fluorescence intensity distributions were analyzed with CytExpert software (Beckman Coulter).

### Statistical analysis

Statistical analyses were performed with GraphPad Prism 8.1.2. Data are presented as mean ± SEM, and n denotes the number of animals unless otherwise stated in the figure legend. Differences between two groups were assessed with the Mann–Whitney test or, where indicated in the figure legends, an unpaired t test. Multiple comparisons were analyzed by Kruskal–Wallis test followed by Dunn’s multiple comparison test, by one-way ANOVA followed by Dunnett’s multiple comparison test, or by two-way ANOVA followed by Sidak’s multiple comparison test; the test applied to each dataset is specified in the corresponding figure legend. Hashtags denote differences relative to the sham + NC5 group and stars denote differences relative to the cancer+NC5 group: p < 0.05 (#/*), p < 0.01 (##/**), p < 0.001 (###/***) and p < 0.0001 (####/****).

## Author contributions

E.M. designed the project, carried out the experiments, analyzed the results and wrote the manuscript, with critical revisions provided by P.S., S.W.K. and J.C. A.T. performed nerve immunofluorescence and analyzed the results. E.M. and J.C. performed the PET and CT acquisitions, and E.M. analyzed the PET and CT data. A.M.J., S.D.R. and M.A.B. synthesized the DsiRNAs. P.S. acquired funding. All authors approved the final version of the manuscript.

## Supporting information

SFig. 1

SFig. 2

SFig. 3

SFig. 4

SFig. 5

SFig. 6

SFig. 7

SFig. 8

SFig. 9

## Acknowledgment

We thank Étienne Croteau and Otman Sarrhini for their help with, and advice on, PET/CT image processing and analysis. Figures were generated with BioRender.com.

