## Supplementary files for "CCR2 silencing in sensory neurons blocks bone cancer progression"

### Supplementary Table Legends

**Supplementary Table 1. Nucleotide sequences of the sense and antisense strands of anti-rCCR2 DsiRNAs**. S3AS12 DsiRNAs were modified by adding 2'-OMethyl-modified RNA bases (m) to the sense and antisense strands.

**Supplementary Table 2.** Sequences of primers used for quantitative real-time RT–qPCR analysis.

### Supplementary Figure Legends

**Supplementary Figure 1: *In vivo* tumor size scores.**

(**A**) Bone anatomy. (**B**) Changes in tumor scores were recorded as follows: score 0 – no tumor, score 1 = slight periosteal distention, score 2 – small tumor outgrowth, score 3 – tumor outgrowth invading the epiphysis, score 4 - tumor outgrowth invading the epiphysis and part of the diaphysis, score 5 – maximum ethical tumor growth prior to animal sacrifice. The corresponding tumor sizes are shown. The higher the score is, the larger the tumor.

**Supplementary Figure 2. Validation of CCR2 silencing and *in vivo* efficacy.**

(**A**) The mRNA interference efficacy of two DsiRNA sequences targeting the CCR2 receptor, CCR2-5U and CCR2-6U (unmodified), was compared to that of the HPRT control gene and to that of their modified (methylated) counterparts CCR2-5 and CCR2-6 before encapsulation into lipid nanoparticles (LNP-DsiRNA). (**B**) Effect of CCL2 (1 µg/rat, i.t.) and coinjection of CCL2 with DsiRNA (CCR2-6 sequence) on the mechanical threshold compared to that of rats receiving NC5 scrambled control DsiRNA. (**C**) Area under the curve of DsiRNA efficacy in reversing CCL2-induced pain responses. (**D**) CCR2 expression and (**E**) p-ERK immunoreactivity in the dorsal root ganglia on day 14 in the sham+NC5, cancer+NC5 and cancer+DsiRNA groups ($n = 3$ to 4 animals per group, 10 sections per animal). The scale bar corresponds to 20 µm. The data are presented as the means ± SEMs. Two-way ANOVA followed by Sidak's multiple comparisons test in (**B**). Kruskal–Wallis test followed by Dunn's multiple comparison test in (**C–E**). $^*P < 0.05$, $^{**}P < 0.01$, $^{***}P < 0.001$. # compared with saline and * compared with CCL2+NC5.

**Supplementary Figure 3: LNPs-DsiRNAs do not alter contralateral bone marrow metabolism, bone density, bone vascularization or animal weight gain.**

(**A**) Repeated daily administrations of LNP-DsiRNA (297 pmol/rat; i.t.). between days 11 and 14 on $^{18}$F-FDG (glucose metabolism), $^{18}$F-NaF (bone structure) and $^{64}$Cu-RGD (blood vessels) mean standardized uptake values (SUV mean) of the whole non-tumor-bearing contralateral femur (cancer+NC5 $n = 5$ and cancer+DsiRNA $n = 6$) and (**B**) animal weight gain (sham+NC5 $n = 7$, cancer+NC5 $n = 15$, cancer+DsiRNA $n = 18$). The data are presented as the means ± SEMs. Mann–Whitney test in (**A, B** and **C**). Two-way ANOVA followed by Sidak's multiple comparisons test in (**D**).

**Supplementary Figure 4: LNPs-DsiRNAs targeting CCR2 decrease substance P expression in DRG neurons induced by bone cancer pain.**

Effect of LNP-DsiRNA on (**A**) substance P and (**B**) CGRP expression in DRG neurons at POD 14. ($n$ = 3 to 4 animals per group, 10 sections analyzed per animal). The scale bar corresponds to 40 µm. The data are presented as the means ± SEMs. Kruskal–Wallis test followed by Dunn's multiple comparisons test in (**A** and **B**). $^*P < 0.05$, $^{**}P < 0.01$, $^{***}P < 0.001$. # compared with the sham+NC5 group and * compared with the cancer+NC5 group.

**Supplementary Figure 5: $^{64}$Cu-RGD radionuclide synthesis and validation.**

(**A**) The $^{64}$Cu-RGD radionuclide structure used in the present study. (**B**) Mass spectrometry and (**C**) ultra-performance liquid chromatography (UPLC) spectra of the $^{64}$Cu-RGD radionuclide. $^{64}$Cu-RGD radionuclide uptake in the (**D**) ipsilateral and contralateral (**E**) bones of animals previously blocked or not blocked with cold RGD peptide (naïve unblocked $n$ = 4 and naïve blocked $n$ = 3). The data are presented as the means ± SEMs. Two-way ANOVA followed by Sidak's multiple comparisons test in (**D** and **E**). $^*P < 0.05$, $^{**}P < 0.01$, $^{***}P < 0.001$. * compared with unblocked cancer animals.

**Supplementary Figure 6: Ziconotide inhibits bone cancer pain-related behaviors but does not decrease tumor growth.**

Effect of twice daily injections of ziconotide (0.5 µg/kg; i.t.) from PODs 11 to 14 on (**A**) mechanical thresholds and (**B**) paw retroflexion scores compared to those of rats receiving saline. (**C**) Representative photographs of the ipsilateral hindpaws of saline- and ziconotide-treated animals and (**D**) femur weights at POD 14 (cancer+saline, $n$ = 6; cancer+ziconotide, $n$ = 7). The data are presented as the means ± SEMs. Two-way ANOVA followed by Sidak's multiple comparisons test in (**A** and **B**). Mann–Whitney test in (**D**). $^*P < 0.05$, $^{**}P < 0.01$, $^{***}P < 0.001$. * compared with the cancer+saline group.

**Supplementary Figure 7: MRMT-1 cancer cells do not express the CGRP or substance P receptors.**

RT–qPCR followed by capillary electrophoresis analyses of the mRNA expression of the CGRP receptor subunits (**A**) CRLR and (**B**) Ramp1 and the substance P receptor (**C**) NK1R in MRMT-1 cells (performed in triplicate).

### Supplementary Materials and Methods

#### Cell culture (Supplementary Fig. 2)

HEK293 and HEKCCR2 cells were harvested in DMEM supplemented with 10% FBS and 2% penicillin/streptomycin. The HEK293 medium also contained 20 mM HEPES. The cells were maintained at 37°C and 5% $CO_2$ in a humidified atmosphere.

#### Immunostaining (Supplementary Fig. 2)

After perfusion, the L1 to L3 DRG were collected, postfixed in 4% paraformaldehyde solution at 4°C for 24 h and then cryoprotected in 30% sucrose in PBS at 4°C for 48 h. Frozen tissues were embedded at −35°C in O.C.T. compound; 30 µm transverse spinal cord sections were generated using a Leica SM220R sliding microtome, and 20 µm DRG sections were generated using a cryostat on gelatin-coated SuperFrost Plus slides. The sections were blocked (10% NGS, 1% BSA, 0.05% Tween-20, 0.1% Triton X-100 in PBS) and incubated in 0.3 M glycine containing 0.2% Tween 20. The sections were labeled with mouse anti-pERK (1:100, Cell Signaling #9101), chicken anti-CCR2 (1:200, Aves Labs, custom) and anti-CGRP (1:500, Millipore #AB5705) in blocking buffer for 24 hours at R.T. The sections were further incubated with fluorophore-conjugated secondary antibodies (1:500, AlexaFluor 488, 568, Invitrogen), and spinal cord sections were mounted on SuperFrost Plus slides. All slides were coverslipped with ProLong Diamond Mountant. $pERK^+$ and $CCR2^+$ neurons were manually counted using ImageJ software. Fluorescence images of the spinal cord and DRG slices were acquired at 20× using a Leica DM4000 microscope equipped with a Leica DFC350FX camera using the same acquisition parameters. Quantification was performed using ImageJ software. Eight to ten slices were analyzed per animal (n=4 sham+NC5, n=4 cancer+NC5, n=3). Representative fluorescence images were acquired using an Olympus FV1000 confocal microscope. All antibodies were validated with positive and negative tissues.

#### *In vitro* DsiRNA transfection and quantitative real-time PCR (RT–qPCR) (Supplementary Fig. 2)

HEK293 cells stably expressing the rat CCR2 receptor (kindly provided by Pfizer, UK) were cultured ($1.3 \times 10^5$ cells) in 48-well culture plates with F12 medium supplemented with 10% fetal bovine serum, 1% penicillin/streptomycin, 0.1 mM nonessential amino acids, 0.75 mg/ml G418, and 1 μg/ml puromycin. When the cells reached 40–50% confluence, three different wells were independently transfected with anti-rCCR2 DsiRNA or the mismatch control (**NC1M7**) at 0.1 nM, 1 nM, or 10 nM and mixed with 10 nM of 1 μL of RNAiMAX (Invitrogen, Carlsbad, CA) 20 min before transfection. Total RNA was extracted 24 h after transfection using an SV96 total RNA isolation system (Promega). RNA quality was verified using a Bioanalyzer 2100 (Agilent). Reverse transcription was performed using 150 ng of total RNA and 20 U of SuperScript II Reverse Transcriptase (Invitrogen) with both random hexamer and oligo-dT primers. Real-time PCR was performed in triplicate for each cDNA sample using Light Cycler 480 probe master mix and a Light Cycler 480 unit (Roche Applied Science). rCCR2 expression levels were analyzed by absolute expression and normalized against the internal controls (the reporter genes hypoxanthine-guanine phosphoribosyltransferase (HPRT) and ornithine decarboxylase (Odc)). The HPRT gene was used as a positive control. The normalized mRNA expression level of rCCR2 obtained from the mismatch negative control group was set at 100%. The sequences of primers used are listed in Table 2.

**CCL2 induced acute pain (Supplementary Fig. 2)**

Animals were randomly assigned to experimental groups and injected intrathecally with saline, CCL2 (0.1 μg/rat, dissolved in 0.1% BSA in sterile saline; R&D Systems #3144-JE-025/CF), CCL2 + NC5 or CCL2 + DsiRNA (297 pmol/rat) under light isoflurane anesthesia. For acute pain tests, DsiRNAs were transfected using Transductine (1:15, Integrated DNA Technologies Inc.).

**$^{64}$Cu-RGD peptide validation (Supplementary Fig. 5)**

Positron emission tomography (PET) imaging was performed using a Triumph™ PET/CT dual modality imaging platform (Gamma Medica, Inc.) equipped with an avalanche photodiode-based digital PET scanner with a 7.5 cm axial field of view. The scanner achieved a transaxial spatial resolution of 1.2 mm and a detection efficiency of 2.6% with an energy window setting of 250-650 keV. A blocking experiment was performed by injecting approximately 35 MBq of $^{64}$Cu-RGD (unblocked) or $^{64}$Cu-RGD coinjected with RGD (blocked), and the accumulation of the radiotracer in the femur was monitored by 30-min dynamic imaging. After each scan sequence, a cylindrical phantom containing a known quantity of free radiotracer, according to the approximate size of the rat (~30 mL), was used to obtain a calibration factor to convert the counts per second into absolute activity measurements in mBq, from which SUVmean and SUVmax values were derived. A density of 1 g/cc was used to convert the fractional uptake per volume into SUVmean and SUVmax values. Images were reconstructed using the Triumph PET/CT software implemented with a 3D-MLEM algorithm using 20 iterations, a span of 63, and a field of view of 80 mm, with a final matrix resolution of 160 × 160 × 128 and a voxel size of 0.5 × 0.5 × 0.597 $mm^3$. CT acquisition was performed over 360° using 512 projections, with a time frame of approximately 250 ms per projection, a field of view of 84.57 mm and a source peak voltage of 60 kVp at 230 µA and 2x2 pixel binning. The single frame was reconstructed in 0.165 × 0.165 × 0.165 $mm^3$ voxels. CT image reconstruction was performed as previously described [59]. PET images were coregistered to the animal's CT image with Amid. Regions of interest (ROIs) were delimited on CT images of the cancer-implanted paw or on the contralateral paw, and radionuclide uptake was quantified using Amid.

**Ziconotide administration**

Ziconotide (Cayman Chemicals) was administered intrathecally at a dose of 0.5 µg/kg to cancer-bearing animals from post implantation days 11 to 14. This dose was selected based on previous studies performed by our group that revealed side effects at higher doses. Since CCR2 DsiRNA silences CCR2 signaling over an extended period, we administered ziconotide twice daily to provide prolonged pain relief.
