## Supplementary figures and images for "CCR2 silencing in sensory neurons blocks bone cancer progression"

### SFig. 1

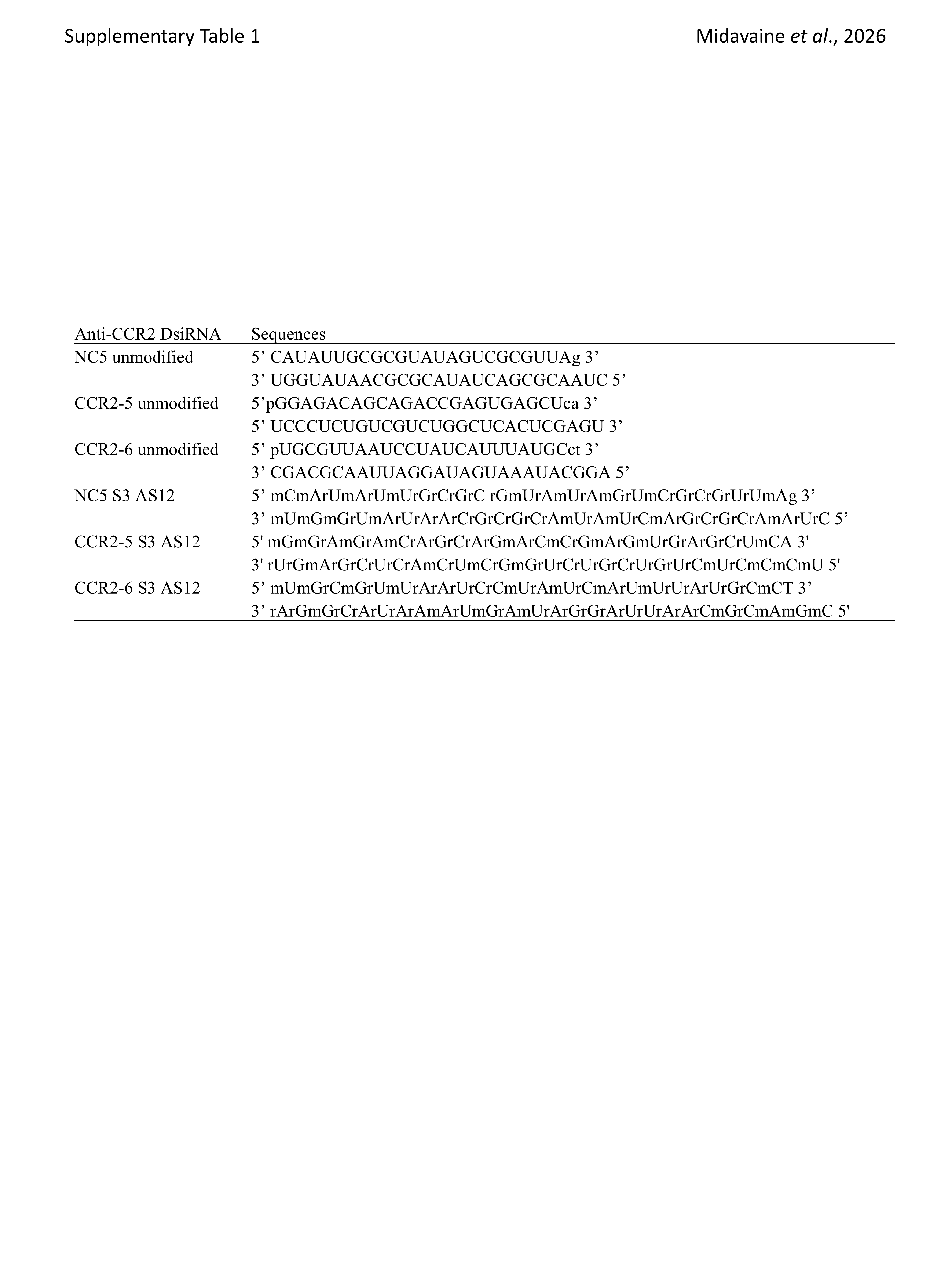

### SFig. 2

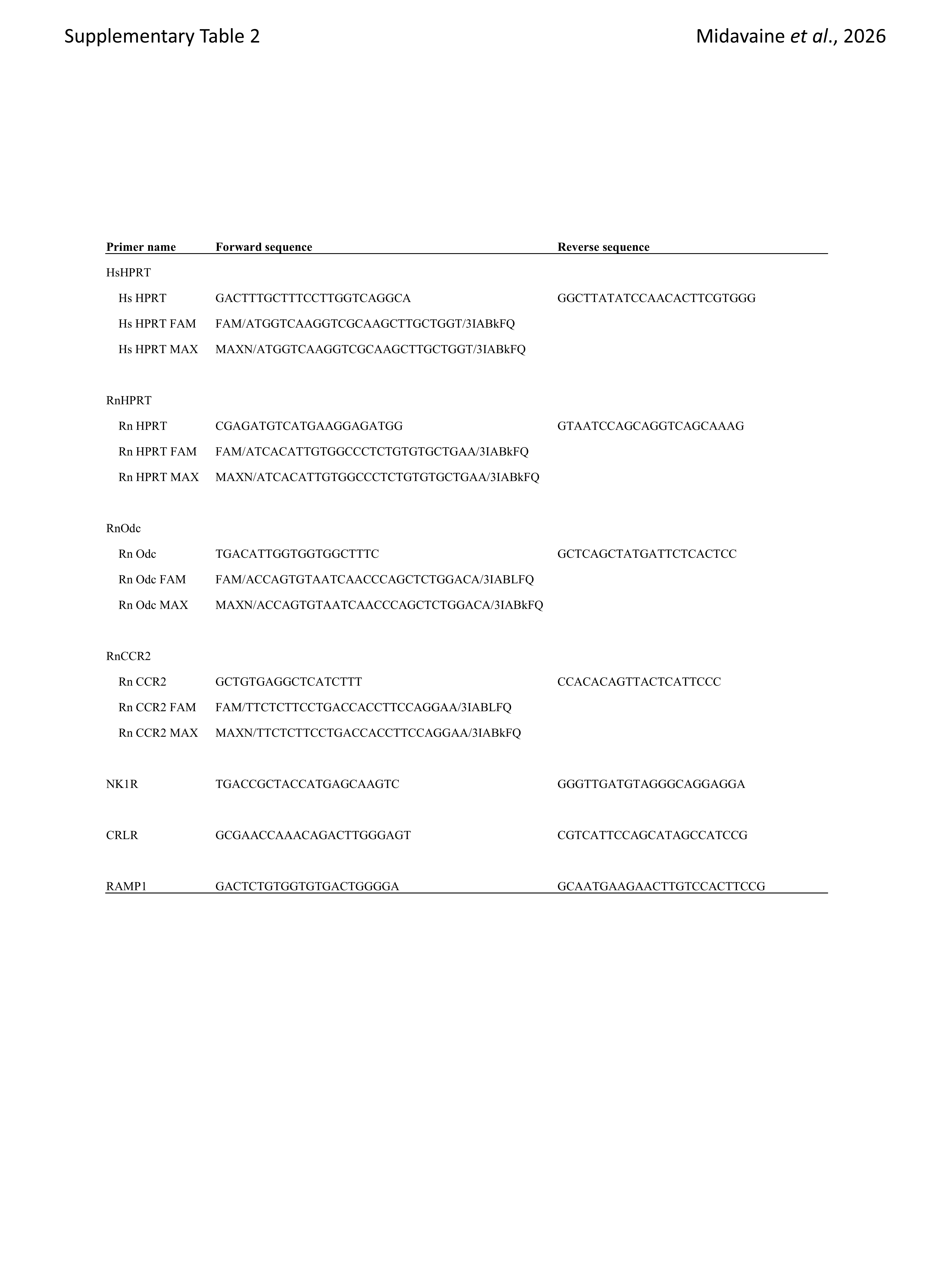

### SFig. 3

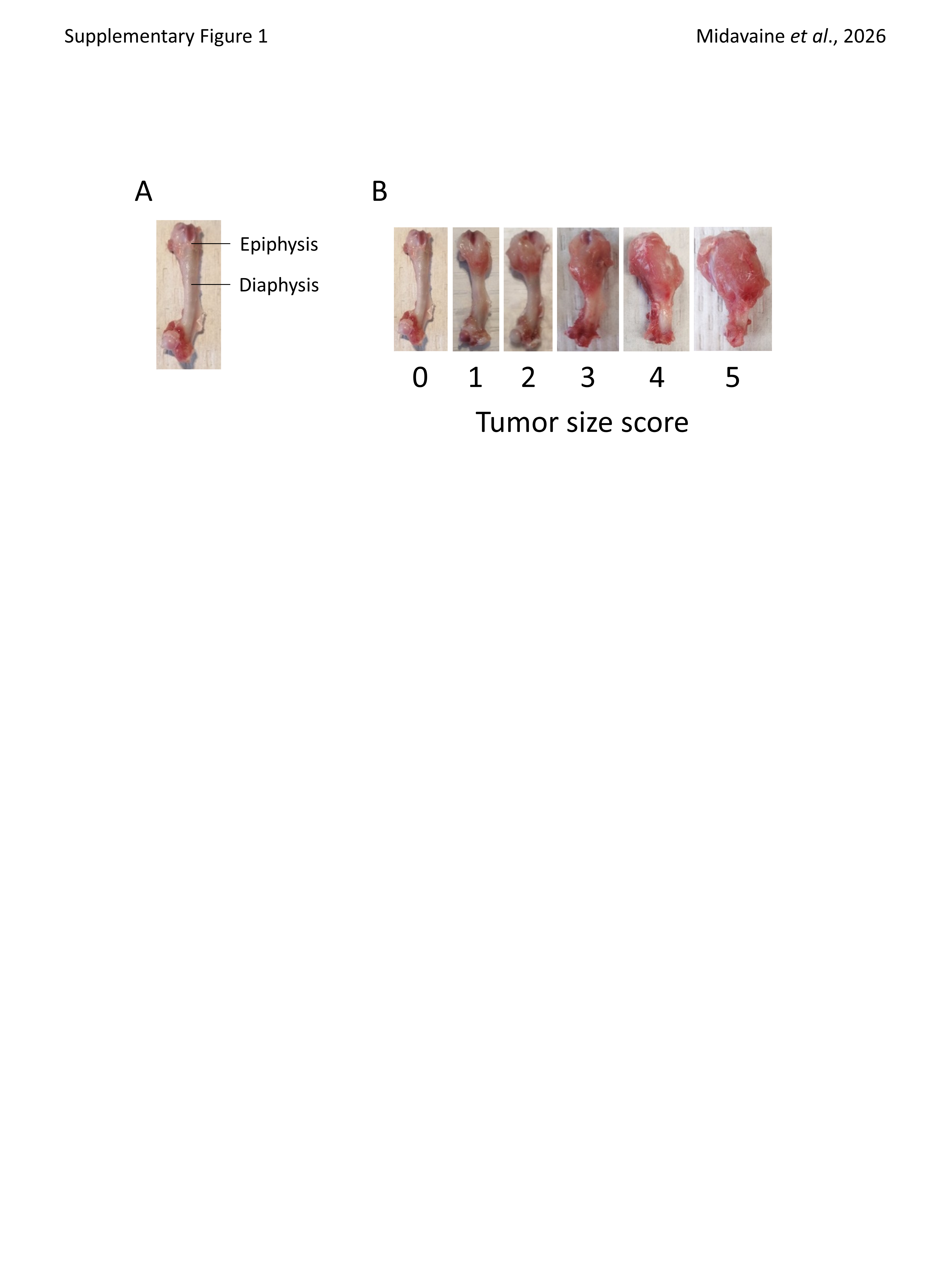

### SFig. 4

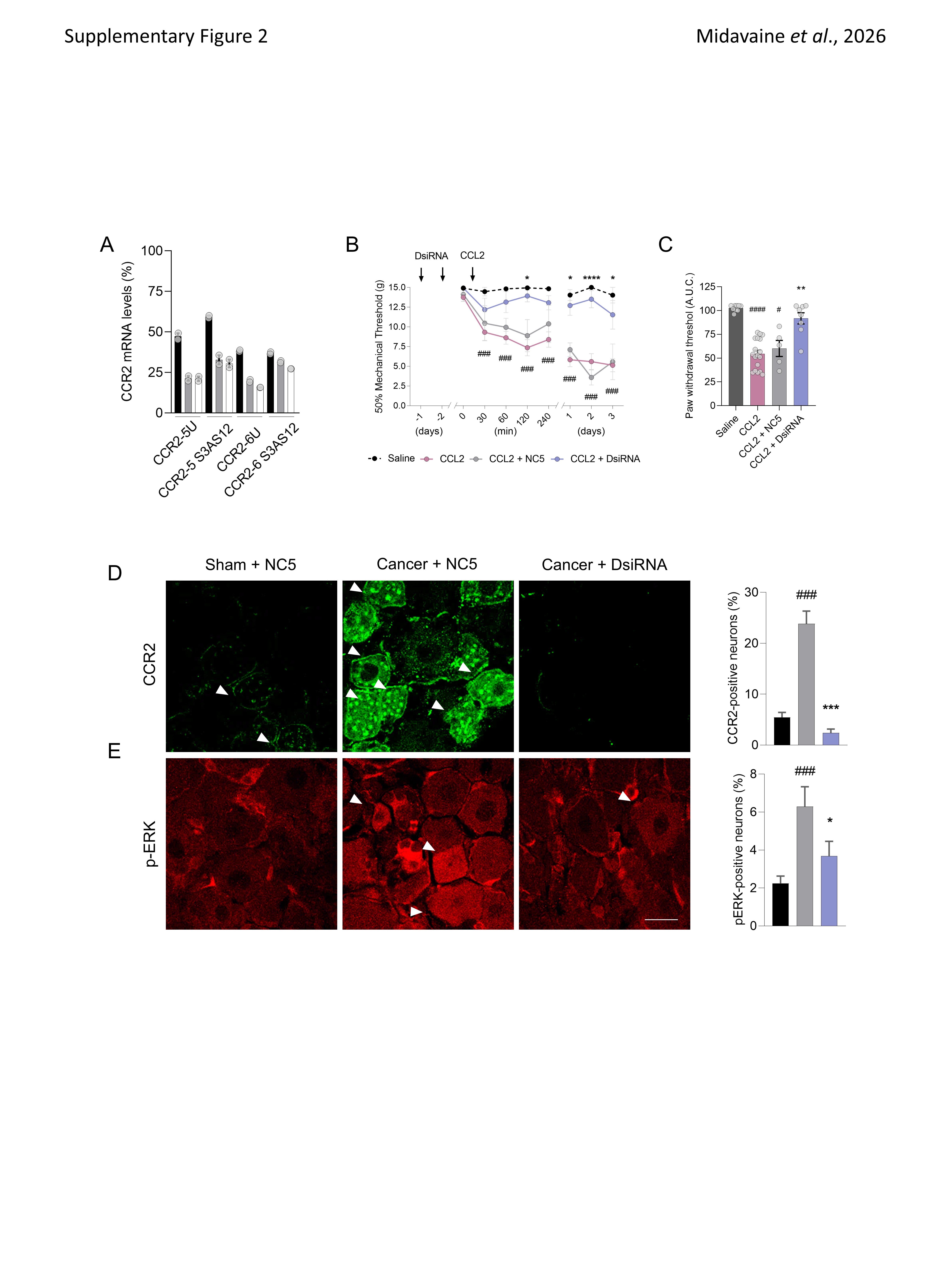

### SFig. 5

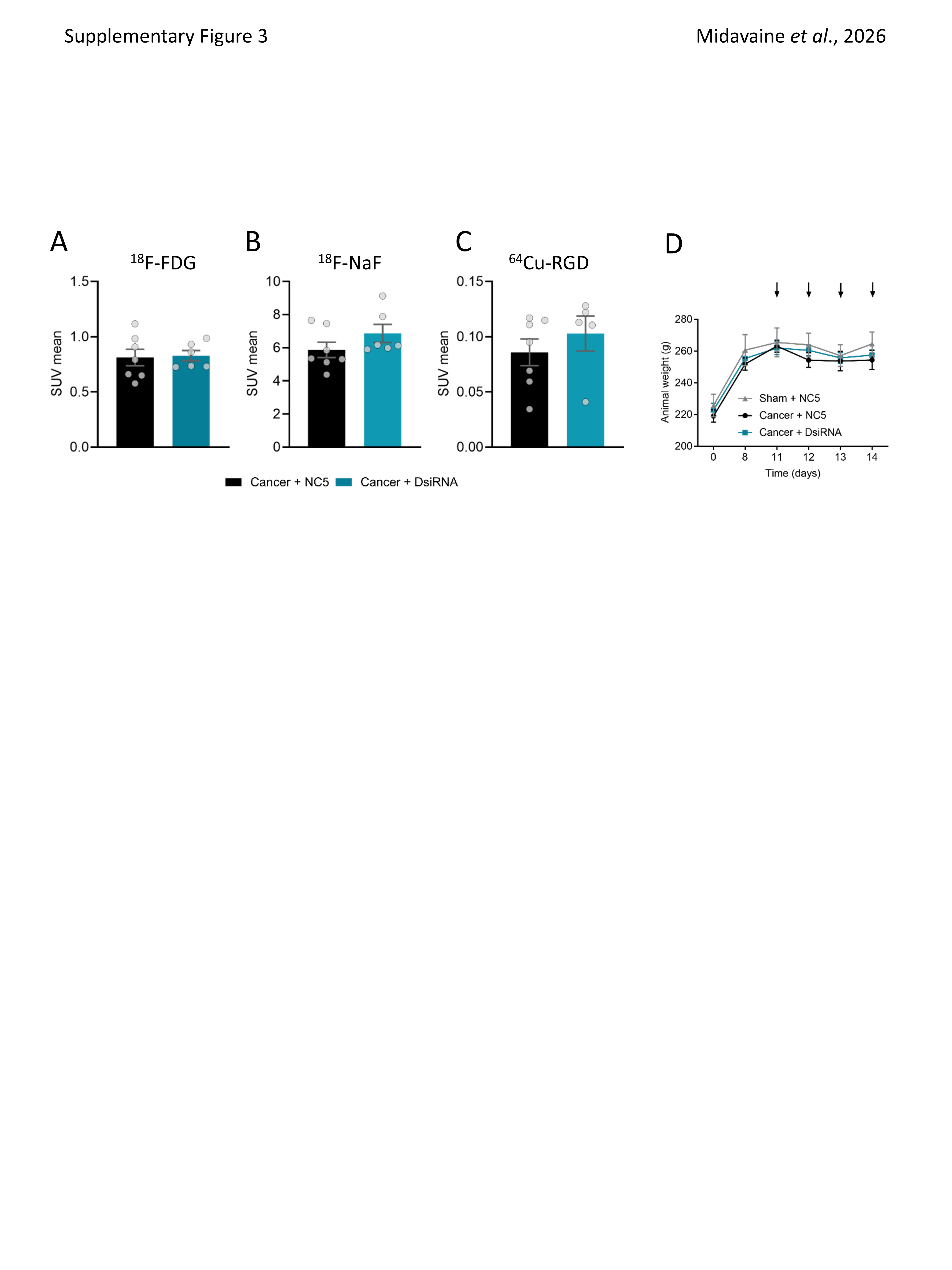

### SFig. 6

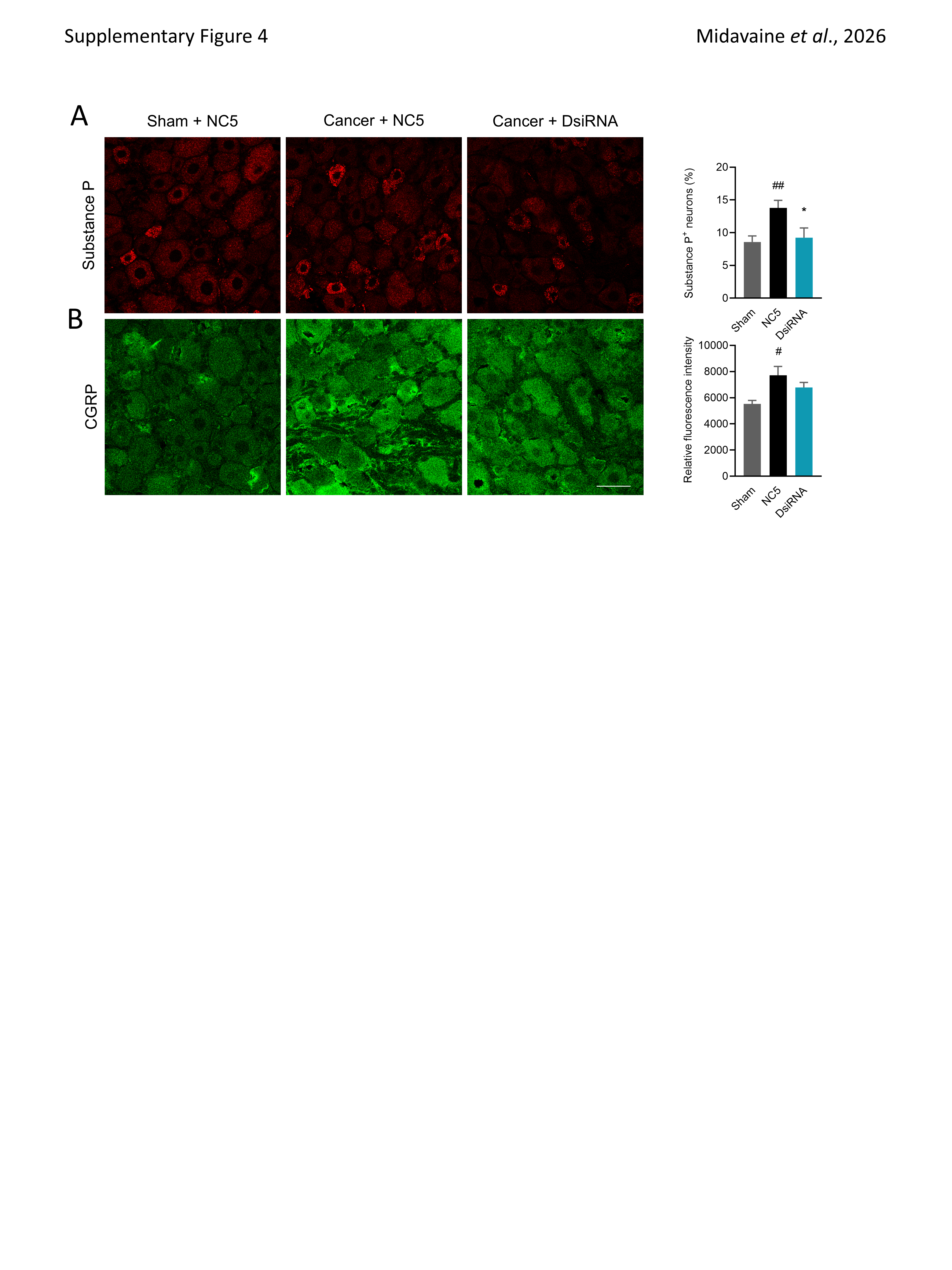

### SFig. 7

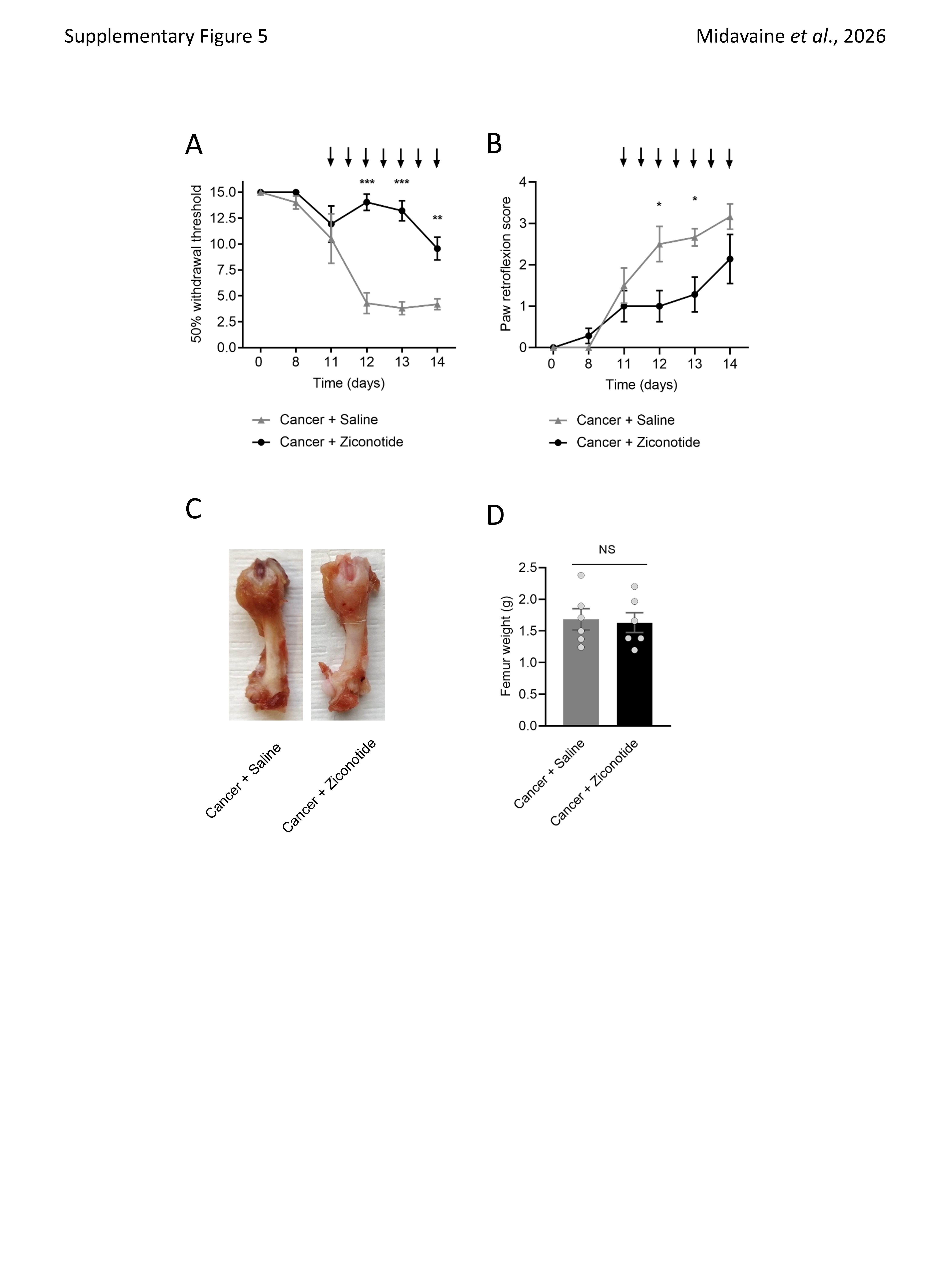

### SFig. 8

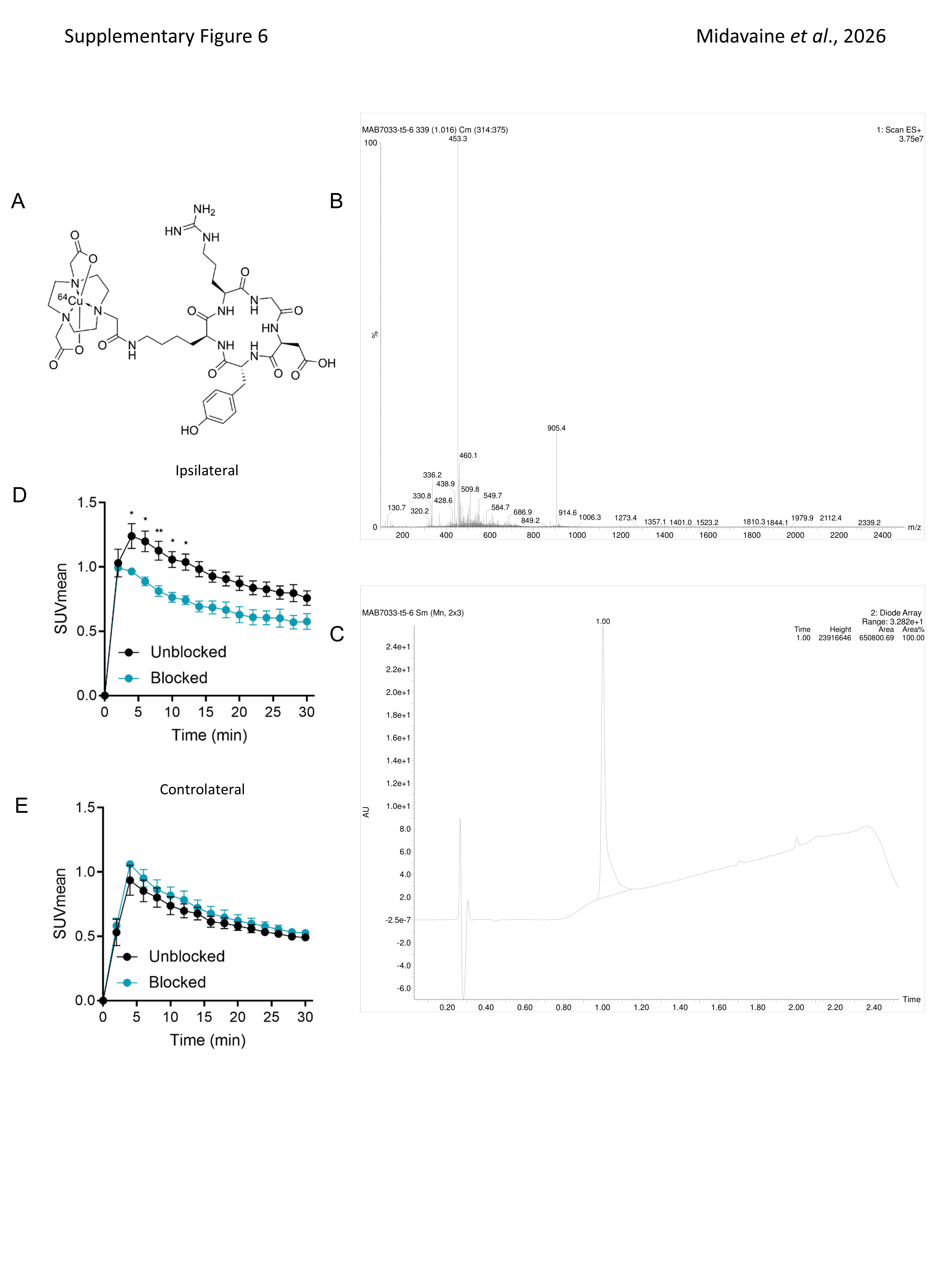

### SFig. 9

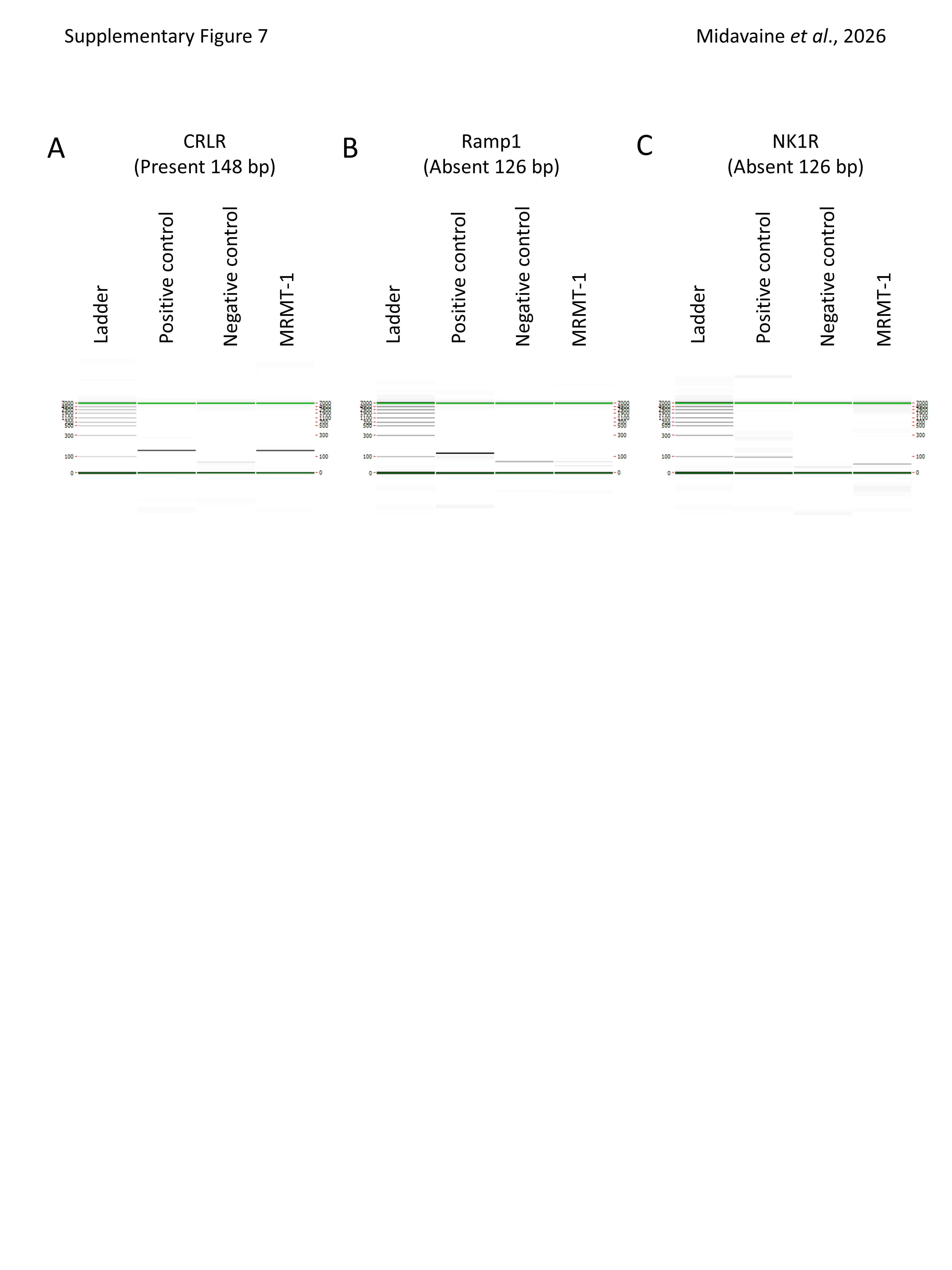
